# Plasticity of Neural Connections Underlying Oxytocin-mediated Parental Behaviors of Male Mice

**DOI:** 10.1101/2021.10.20.465207

**Authors:** Kengo Inada, Mitsue Hagihara, Kazuko Tsujimoto, Takaya Abe, Ayumu Konno, Hirokazu Hirai, Hiroshi Kiyonari, Kazunari Miyamichi

**Author notes:** Correspondence (K.I.), (K.M.).

## Abstract

The adult brain can flexibly adapt behaviors to specific life-stage demands. For example, while sexually naïve male mice are aggressive to the conspecific young, they start to provide caregiving to infants around the time when their own young are expected. How such behavioral plasticity is implemented at the level of neural connections remains poorly understood. Using viral-genetic approaches, here we establish hypothalamic oxytocin neurons as key regulators of parental caregiving behaviors of male mice. We then used rabies virus-mediated unbiased screen to identify excitatory neural connections originating from the lateral hypothalamus to the oxytocin neurons to be drastically strengthened when male mice become fathers. These connections are functionally relevant, as their activation suppresses pup-directed aggression in virgin males. These results demonstrate the life-stage associated, long-distance, and cell-type-specific plasticity of neural connections in the hypothalamus, the brain region classically assumed to be hard-wired.

**Highlight:**

– OT is indispensable for parental caregiving behavior of male mice
– Activation of OT neurons triggers paternal caregiving behavior in otherwise infanticidal sexually-naïve male mice partly via OT ligand
– Unbiased rabies virus-mediated screening reveals enhanced connectivity originated from excitatory LHA neurons to OT neurons in fathers.
– This structural plasticity can support behavioral plasticity

## Introduction

The adult brain is equipped with the plasticity to change behaviors to specific life-stage demands. For instance, sexually-naïve animals rarely take care of the conspecific young, as caregiving behaviors can bring heavy burdens and may decrease their fitness (Dulac et al., 2014). The emergence of caregiving behaviors towards infants is broadly observed across the animal kingdoms that occurs around a time when their own young are expected (Elwood, 1994). Despite the accumulating behavioral observations, neural mechanisms that give rise to the behavioral changes remain poorly understood. A major mechanism by which the brain achieves plasticity is through changes in the strength of synaptic connections. However, it is highly challenging to pinpoint which neurons (or neural types) form synapses that undergo plastic change upon any biological event. Solving this issue requires a strategy to identify the precise location and cell types of neurons whose synaptic connections experience plastic change. We herein aim to provide a specific example of this challenge by examining the extent to which cell-type-specific connection diagrams change in the adult male mice when they become fathers.

How are male animals engaged in direct nurturing behaviors to infants? Studies of rodent model animals such as prairie voles, rats, and mice have suggested molecular and neural mechanisms underpinning paternal caregiving behaviors (Dulac et al., 2014; Kohl and Dulac, 2018; Rilling and Young, 2014; Yoshihara et al., 2018). For example, lesion studies have revealed the medial preoptic nucleus (MPN) as a positive regulator of paternal caregiving behaviors (Lee and Brown, 2002; Tsuneoka et al., 2013). Specifically, MPN neurons expressing galanin (Kohl et al., 2018; Wu et al., 2014) and calcitonin receptor (Calcr) (Moffitt et al., 2018; Yoshihara et al., 2021b), whose functions are well documented in maternal behaviors, also contribute to paternal caregiving. In addition, the prolactin signaling in the MPN is required for paternal caregiving (Stagkourakis et al., 2020). On the other hand, male animals are equipped with neural substrates located in multiple brain regions for infant-directed aggression (infanticidal) behaviors (Autry et al., 2021; Chen et al., 2019; Sato et al., 2020; Tsuneoka et al., 2015), which should be suppressed for the execution of paternal care. Yet it remains largely elusive as to how male animals turn to favor caregiving over infanticide upon the life-stage transition.

Neural hormone oxytocin (OT) produced by the OT neurons located in the paraventricular hypothalamic nucleus (PVH) has received attention in studies of maternal caregiving behaviors (Rilling and Young, 2014). Intracerebroventricular (icv) (Pedersen et al., 1982) or intraperitoneal (ip) (Marlin et al., 2015) injection of OT induces caregiving behaviors in virgin rodent females, and so does optogenetic activation of the OT neurons in the PVH (Marlin et al., 2015; Scott et al., 2015). In contrast, loss-of-function of OT or its receptor, OTR, shows only minor phenotypes of maternal caregiving behaviors (Insel and Young, 2001; Macbeth et al., 2010; Nishimori et al., 1996; Young et al., 1996). These studies suggest that OT signaling can facilitate the onset, but to a lesser extent the maintenance of, maternal care (Yoshihara et al., 2018). Despite the importance of OT in mothers, OT signaling in the male brain in relation to paternal caregiving behaviors has been overlooked in classical studies of biparental model animals (Rilling and Young, 2014) or genetic mutant mouse studies. As neural hormones are suited for modulating multiple neural circuits across the brain, we envision that OT may play a central role in the behavioral plasticity of male mice when they become fathers.

In the present study, we first establish crucial functions of OT neurons and OT ligands for paternal caregiving behaviors by a series of viral genetic experiments in male mice. We then aim to identify pre-synaptic partners of OT neurons that undergo structural changes in connections upon paternal life-stage transition. For this, we utilize rabies-virus mediated retrograde trans-synaptic tracing (Miyamichi et al., 2013) as an unbiased screening method (Beier et al., 2017; Callaway and Luo, 2015). This led us to identify specific excitatory neural connections to OT neurons that are enhanced in fathers.

## Results

### OT is necessary for the parental caregiving behaviors

We first examined whether OT secretion from OT neurons is necessary for the expression of parental behaviors in fathers. To this end, by CRISPR/Cas9-mediated genome editing, we generated *OT* deletion (−) and floxed (flox) alleles and prepared *OT* conditional knockout (cKO) mice (*flox/*−, Figures 1A and S1A–S1C). Each male mouse was injected with adeno-associated virus (AAV) driving Cre into the PVH, crossed with a female mouse, and co-housed with the partner throughout her pregnancy and parturition. The behavioral assay was conducted five days after the birth of pups (Figure 1B, see Methods). Post-hoc histochemical analyses revealed that injection of *AAV-Cre* deleted floxed *OT* exon 1, resulting in the loss of transcription of *OT* mRNA (Figure 1C). In the behavioral assay, after removing a dam and pups, three unfamiliar pups were introduced into the home cage, and a father mouse was allowed to interact with them freely (see Methods). Unlike control fathers (sham), Cre-injected fathers (+Cre) mostly ignored pups without providing parental caregiving behaviors (Figure 1D and 1F–1I). The latency to investigate the pups was unaffected, implying that their sensation of pups was grossly normal (Figure 1E). The number of remaining *OT*+ neurons, visualized by *in situ* hybridization (ISH), was proportional to the duration of parental behaviors, indicating that OT secretion from a sufficient number of neurons is necessary to exert parental behaviors in fathers (Figure 1H). Similar to the *OT* cKO, whole-body *OT* KO fathers showed defects in the expression of parental behavior (Figure S1D–S1H). These results indicated that OT is indispensable for the expression of parental caregiving behaviors of male mice.

**Figure 1.**
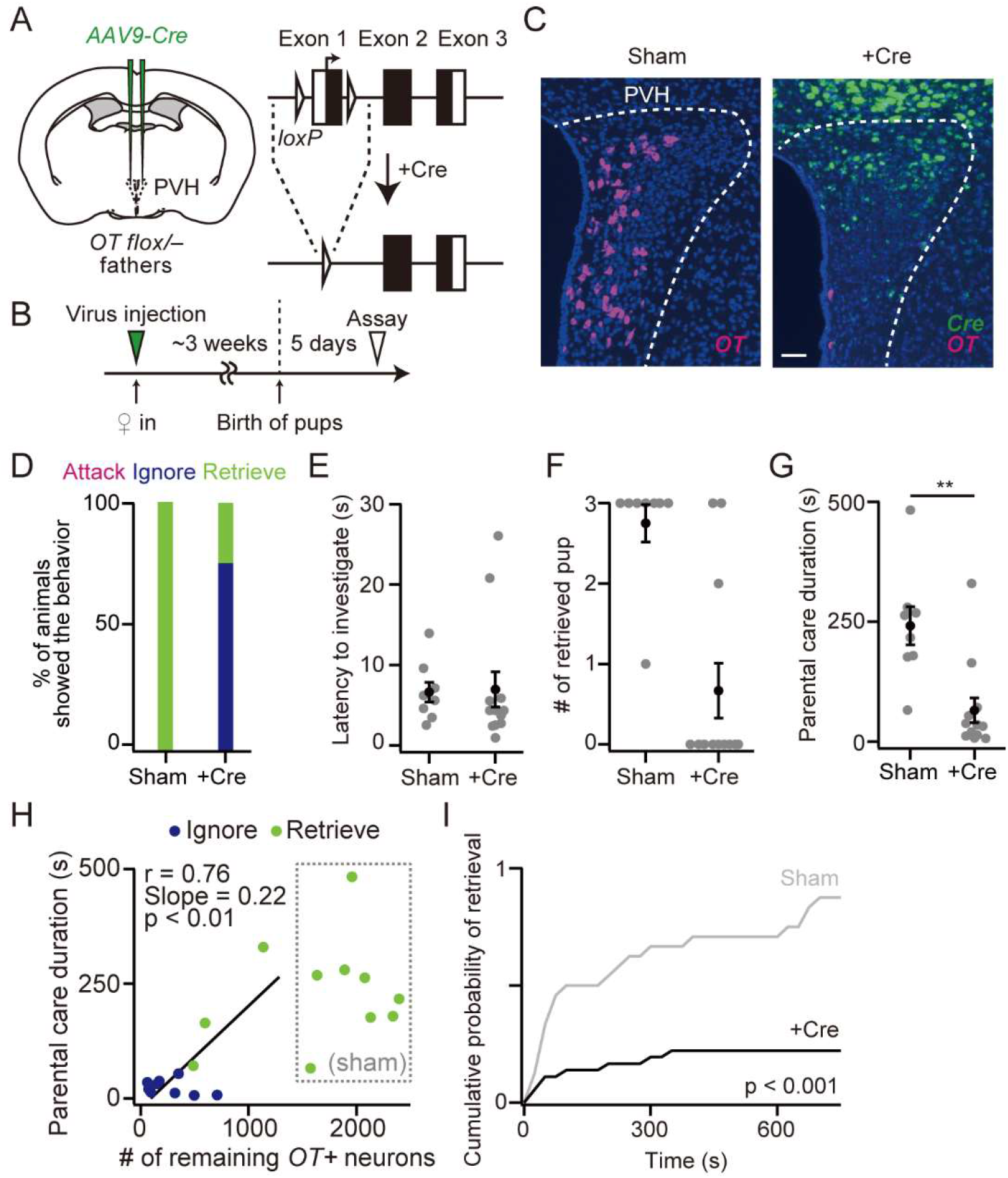
OT is necessary for parental caregiving behaviors. (A) Left: Schematic of the virus injection. *AAV-Cre* was injected into the bilateral PVH of *OT*^*flox*/−^ male mice. Right: Schematic of the Cre expression and excision of floxed exon 1 of the *OT* gene. (B) Schematic of the timeline of the experiment. (C) Representative coronal sections of the PVH from *OT*^*flox*/−^ fathers without (sham; left) or with (+Cre; right) *AAV-Cre* injection. *OT* and *Cre in situ* staining were shown in magenta and green, respectively. Blue, DAPI. Scale bar, 50 μm. (D) Percentage of fathers showing attack, ignore, or retrieve. (E) Latency to the first investigation of pups was not statistically different (p > 0.9, two-tailed Welch’s *t*-test). (F) The number of retrieved pups. (G) Parental care duration. **p = 0.0039, two-tailed Welch’s *t*-test. (H) Correlation between the number of remaining *OT*+ neurons in the PVH and parental interaction. Black line, a linear fit for +Cre fathers. (I) Cumulative probability of pup retrieval. The p-value is shown in the panel (Kolmogorov– Smirnov test). n = 8 and 12 for sham and Cre-injected, respectively. Error bars, standard error of mean (SEM). See Figure S1 for more data.

### OT neurons can evoke caregiving behaviors in virgin males

To examine the gain-of-function effect of OT neurons, we chemogenetically activated OT neurons in PVH by expressing *hM3Dq-mCherry* (Figure 2A and 2B). *Ex vivo* whole-cell patch-clamp recordings from hM3Dq+ neurons confirmed that clozapine N-oxide (CNO) evoked burst firing (Figure S2A and S2B). We ip injected saline or CNO 30 min before the behavioral assay (Figure 2C). CNO injection inhibited pup-directed aggression and significantly facilitated pup retrieval (Figure 2D–2I), which was not observed in mCherry control (Figure S2C–S2J).

**Figure 2.**
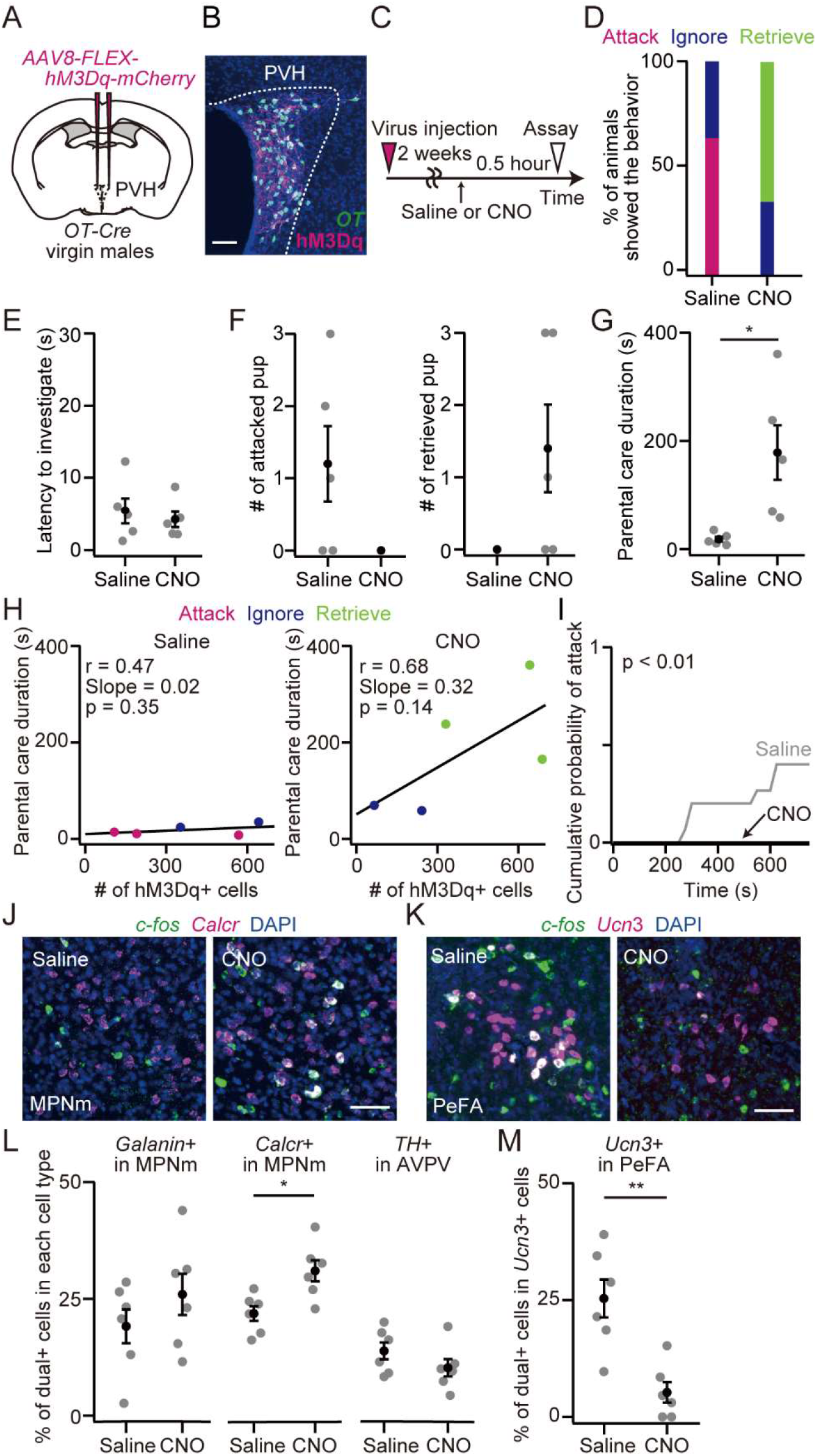
Activation of OT neurons facilitates parental behavior in virgin males. (A) Schematic of the virus injection. *AAV-FLEx-hM3Dq-mCherry* was injected into the bilateral PVH of virgin males. (B) Representative coronal section showing the expression of *OT* mRNA (green) and *hM3Dq-mCherry* (magenta). Blue, DAPI. Scale bar, 50 μm. (C) Schematic of the timeline of the experiment. (D) Percentage of animals showing attack, ignore, or retrieve. In the virgin males that expressed hM3Dq, CNO suppressed pup-directed aggression and promoted parental behavior. n = 6 and 6 for saline and CNO in the hM3Dq, respectively. (E) Latency to the first investigation of pups. (F) Numbers of attacked (left) and retrieved (right) pups. (G) Duration of parental care (*p = 0.0470, two-tailed Welch’s *t*-test). (H) Relationship between parental interaction and the number of hM3Dq+ neurons in the PVH. Animals are color-coded by their behavioral categories. Black line represents a linear fit (p = 0.35 and 0.14 for saline and CNO, respectively). (I) Cumulative probability of attack to pups. The p-value is shown in the panel (Kolmogorov– Smirnov test). (J, K) Representative coronal sections of *OT-Cre* virgin males expressing hM3Dq that interacted with pups. Green and magenta represent *c-fos* mRNA and *Calcr* mRNA in the MPNm (J) or *c-fos* mRNA and *Ucn3* mRNA in PeFA (K) *in situ* staining, respectively. Blue, DAPI. Scale bar, 50 μm. (L) Percentage of activated cells in each cell type associated with parental behavior. *Calcr+* cells in the MPNm were activated by the application of CNO. *p = 0.0138, two-tailed Welch’s *t*-test with Bonferroni correction. n = 6 mice for each condition. (M) Percentage of activated cells in *Ucn3+* cells in the PeFA associated with pup-directed aggression in males. The activity of those cells was suppressed by the chemogenetic activation of PVH OT neurons (**p = 0.0040, two-tailed Welch’s *t*-test). n = 6 and 6 for saline and CNO, respectively. Error bars, SEM. See Figure S2 for more data.

These results led us to hypothesize that chemogenetic activation of OT neurons would (i) activate the brain regions facilitating caregiving behaviors, and (ii) suppress the neural activities in brain regions mediating infanticidal behaviors. To examine these possibilities, we analyzed *c-fos* mRNA expression as a readout of neural activation following the chemogenetic stimulation of OT neurons in virgin males. We found elevated neural activities of *Calcr*+ neurons in the medial part of MPN (MPNm), which is known as a center for parental behaviors (Moffitt et al., 2018; Yoshihara et al., 2021a) (Figure 2J and 2L). By contrast, the neural activities of a known center for infanticide, *urocortin 3* (*Ucn3*)*+* neurons in the perifornical area (PeFA)(Autry et al., 2021), were suppressed (Figure 2K and 2M). These findings indicated that the limbic neural populations directly related to parental and infanticidal behaviors are modulated by OT neurons, thereby enabling the execution of fully parental behaviors in otherwise infanticidal virgin males.

### Non-OT and OT ligand synergistically facilitate caregiving behaviors

OT neurons release not only OT but also other neurotransmitters or neuropeptides. For example, OT neurons are glutamatergic, as they express *vesicular glutamate transporter type 2* (*vGluT2*) (Xu et al., 2020). Our gain-of-function experiments (Figure 2) may be mediated by non-OT neurotransmissions. To examine this scenario, we targeted *hM3Dq-myc* to the PVH of *OT^−/−^* virgin males under the control of a 2.6 kb mouse *OT* promotor (*OTp*), which is orthologous to the established rat *OTp* (Knobloch et al., 2012)(Figure 3A and 3B). We first validated that mouse *OTp* selectively labeled OT neurons in virgin male mice: 91.9 ± 1.02 % of neurons expressing hM3Dq-Myc also expressed *OT* mRNA (Figure 3C). We found that OT neurons persisted in the absence of functional *OT* gene, as the number of hM3Dq-Myc expressing neurons did not differ between *OT*^+/+^ and *OT^−/−^* mice (Figure S3A). We then conducted ip injection of saline or CNO, thirty minutes before the behavioral assay (Figure 3D). In the *OT^−/−^* mice, CNO injection leads to the release of neurotransmitters or neuropeptides other than OT. We found that in the saline-injected controls, the pup-directed attack was similarly observed in *OT*^+/+^ and *OT^−/−^* virgin males (Figure 3E), suggesting that OT is not involved in infanticide. This pup-directed attack was markedly suppressed in CNO-injected mice, regardless of the functional *OT* gene (Figures 3E, 3F, and S3B–S3D). Thus, activation of OT neurons can suppress the pup-directed attack independently from OT ligand. However, the caregiving behaviors induced by CNO injection were more apparent in *OT*^+/+^ than *OT^−/−^* mice, as demonstrated by the better performance (Figure 3G and 3H). In addition, a lower number of hM3Dq+ neurons triggered pup retrieval in *OT*^+/+^ mice (Figure 3I and 3J). These data collectively revealed that the OT ligand promotes caregiving behaviors in virgin males, with additional contributions made by other neurotransmitters in the OT neurons.

**Figure 3.**
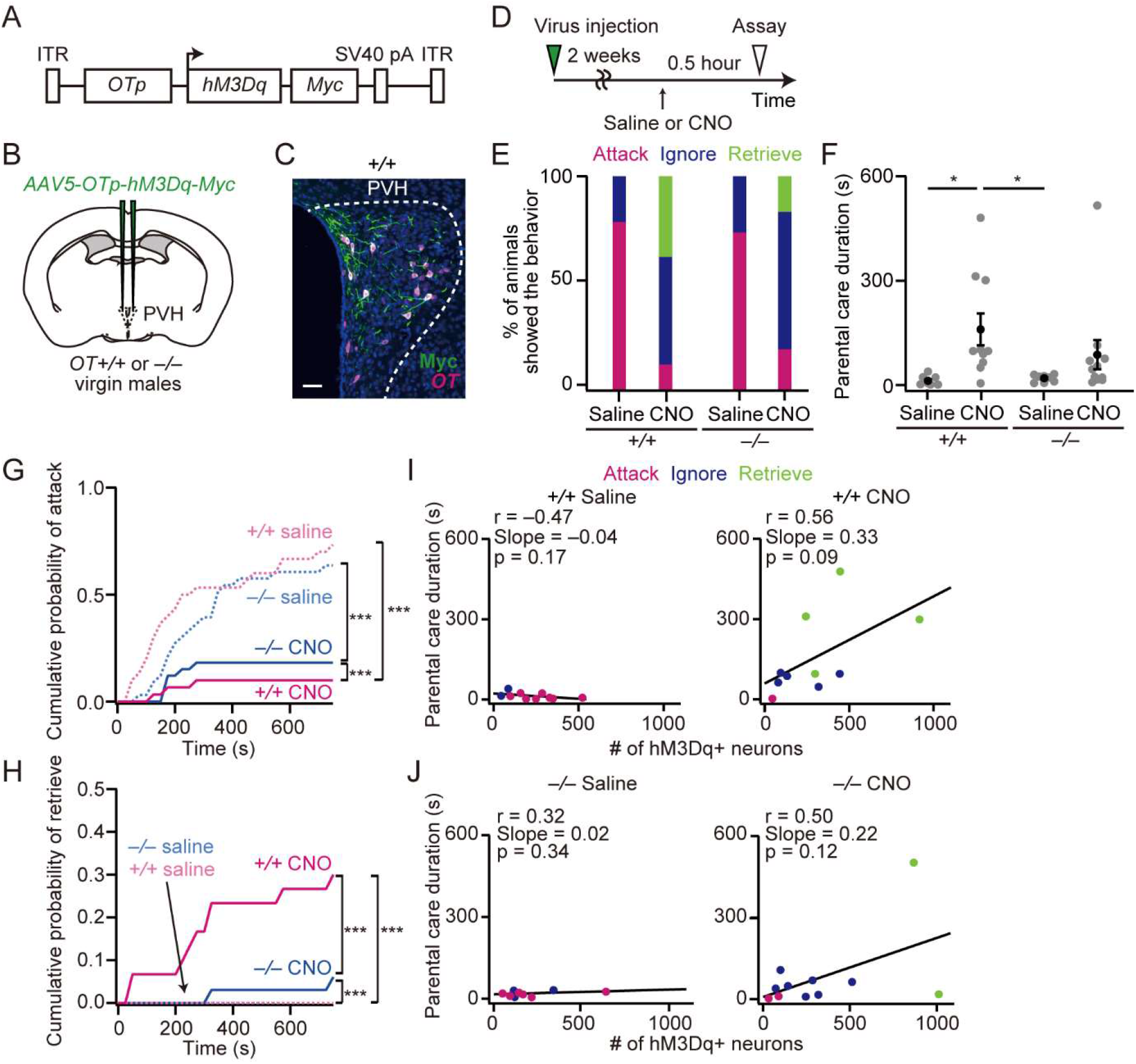
OT ligand promotes the expression of parental behavior in virgin males. (A) AAV construct for driving *hM3Dq-Myc* under a 2.6-kb mouse *OT* promoter (*OTp*). (B) Schematic of the virus injection. (C) Representative coronal sections showing expression of Myc (green) in *OT+* neurons (magenta) with DAPI staining (blue). Scale bar, 50 μm. (D) Schematic of the timeline of the experiment. (E) Percentage of virgin males showing attack, ignore, or retrieve. (F) Parental care duration. *p = 0.0171, *p = 0.0187, one-way ANOVA with post hoc Tukey HSD. (G, H) Cumulative probability of attack (G) or retrieval (H). ***p < 0.001, Kolmogorov–Smirnov test. (I, J) Correlation between the number of hM3Dq+ neurons in the PVH and parental interaction of *OT*^+/+^ (I) or *OT*^−/−^ virgin males (J). Black line, a linear fit. p-values are shown in each panel. Note that retrieval was evoked by a lower number of hM3Dq+ neurons (about 1000 neurons in *OT*^−/−^ and about 400 neurons in *OT*^+/+^). *OT*^+/+^, n = 10 each, and *OT*^−/−^, n = 11 each, for saline and CNO. Error bars, SEM. See Figure S3 for more data.

### Plasticity of monosynaptic connections to PVH OT neurons

Our data thus far have established PVH OT neurons as a crucial hub of caregiving behaviors in male mice. We next aimed to analyze possible changes of neural connections to OT neurons upon life-stage transition. We selectively targeted rabies virus-based retrograde trans-synaptic tracing (Miyamichi et al., 2013) to PVH OT neurons in virgin males and fathers and quantitatively compared the distribution of presynaptic neurons labeled with GFP (Figure 4A). The number of starter cells, defined as the overlap of TVA-mCherry and rabies-derived GFP, was comparable (Figure 4B), implying that the population of OT neurons was kept grossly unchanged from virgin males to fathers. OT neurons received predominant inputs from various hypothalamic nuclei (Figure 4C and 4D) and the input fraction was largely similar between virgin males and fathers (Figure 4D). Closer investigation of the convergence index, defined by the number of labeled presynaptic neurons normalized to the number of starter cells, revealed that the lateral hypothalamus (LHA) and MPNm exhibited more viral labeling in fathers than in virgin males (Figure 4E and 4F), suggesting stronger connections to OT neurons in fathers.

**Figure 4.**
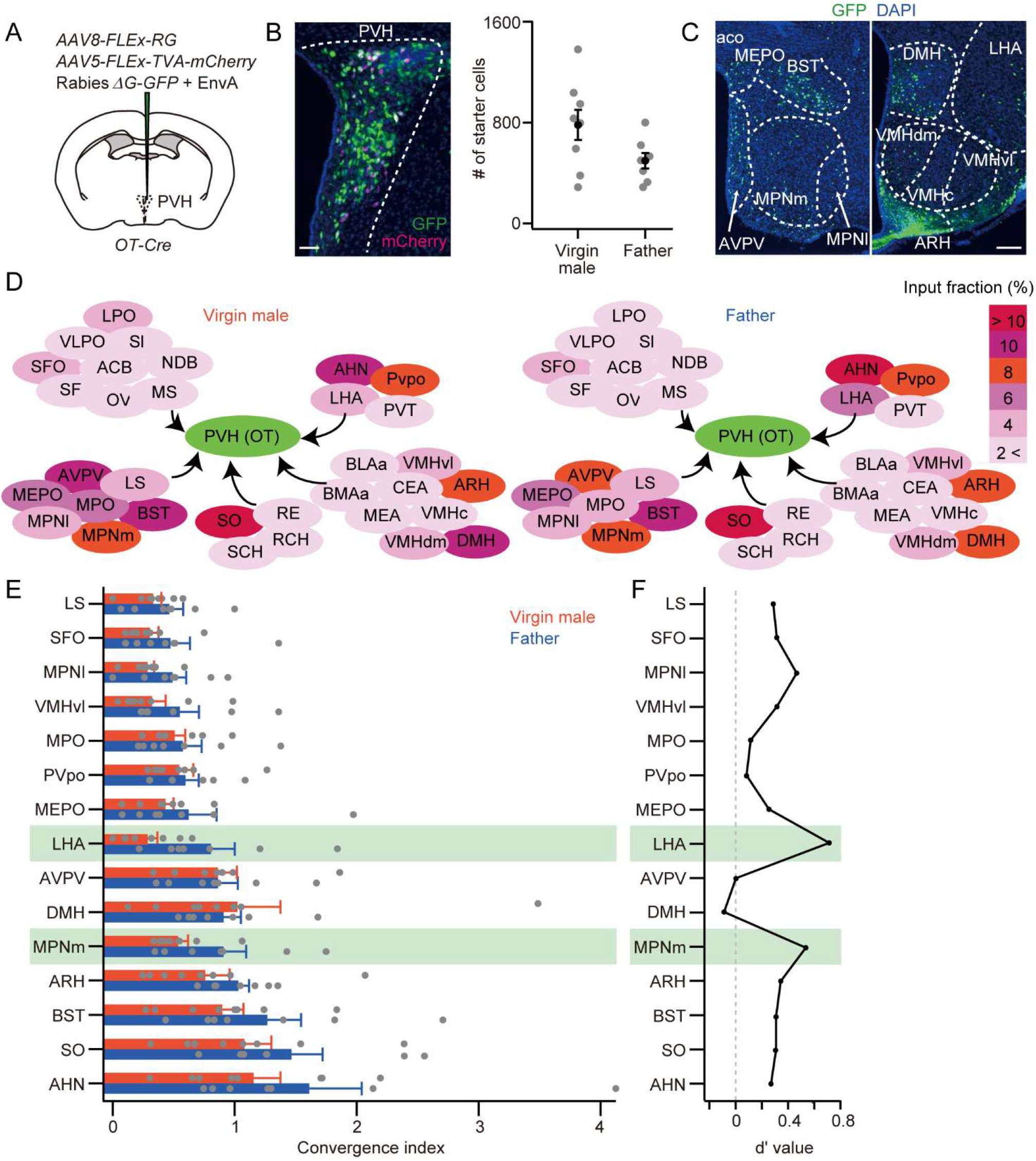
Identification of monosynaptic connections to PVH OT neurons in virgin males and fathers. (A) Schematic of the virus injections. (B) Left: Representative coronal section containing starter cells (white) defined as the overlap of mCherry (magenta) and GFP (green). Blue, DAPI. Scale bar, 50 μm. Right: Number of starter cells in the PVH in virgin males and fathers (p = 0.0751, two-tailed Welch’s *t*-test). (C) Representative coronal sections of presynaptic cells revealed by rabies-GFP. Scale bar, 100 μm. See Table S1 for abbreviations. (D) Fraction of monosynaptic inputs to PVH OT neurons in virgin males (left) and fathers (right). The input pattern was largely invariant. See Table S2 for the complete data set. (E) Normalized inputs from the top 15 nuclei to the OT neurons in virgin males and fathers. See Table S3 for the complete data set. (F) Quantification of the difference of convergence index shown in (E) by d′ value. A d′ value larger than 0 represents a single PVH OT neuron in fathers receiving denser input. A larger d′ value was found in the input from LHA and MPNm, marked by green shadow. n = 8 and 7 for virgin males and fathers, respectively. Error bars, SEM. See Figure S4 for more data.

To test whether the enhanced rabies labeling observed in fathers persists or not, we utilized trans-synaptic tracing in fathers that did not contact pups for 5 weeks (Figure S4A). In both LHA and MPNm, the increased viral labeling observed in 5 days after the birth of pups was returned to a similar level as virgin males after 5 weeks of isolation (Figure S4B–S4F), suggesting that this connection enhancement is transient and reversible.

Altered neural connections may also occur inside the PVH. To examine this possibility, we utilized a modified version of rabies tracing suited for local circuit mapping (Miyamichi et al., 2013) (Figure S4G). PVH contains four major cell types, OT neurons, vasotocin (VT, previously known as arginine vasopressin) neurons (Theofanopoulou et al., 2021), corticotropin-releasing hormone (CRH) neurons, and thyrotropin-releasing hormone (TRH) neurons (Figure S4H). Although we identified VT and CRH neurons as prominent presynaptic partners of OT neurons within the PVH, no major difference was found in the input patterns to OT neurons within the PVH between virgin males and fathers (Figure S4I and S4J).

Next, we identified the cell types of presynaptic neurons at long-distance by histochemical methods. First, we classified them into two types: *vGluT2*+ excitatory neurons and *vesicular γ-aminobutyric acid transporter* (*vGAT*)+ inhibitory neurons. The ratio of excitatory and inhibitory input was largely unchanged, with several exceptions (Figure 5A). The product of convergence index (Figure 4E) and excitatory/inhibitory ratio (Figure 5A) revealed that particularly LHA and MPNm send increased excitatory drive to OT neurons in fathers (Figure 5B). Further analysis of cell-type-specific marker genes (Mickelsen et al., 2019; Moffitt et al., 2018) showed a selective enhancement of input from melanin-concentrating hormone (MCH)-producing neurons in LHA in fathers (Figure 5C–5F). Collectively, rabies-mediated tracing revealed the presynaptic landscape of OT neurons and highlighted excitatory inputs from LHA and MPNm as candidate connections that are strengthened in fathers.

**Figure 5.**
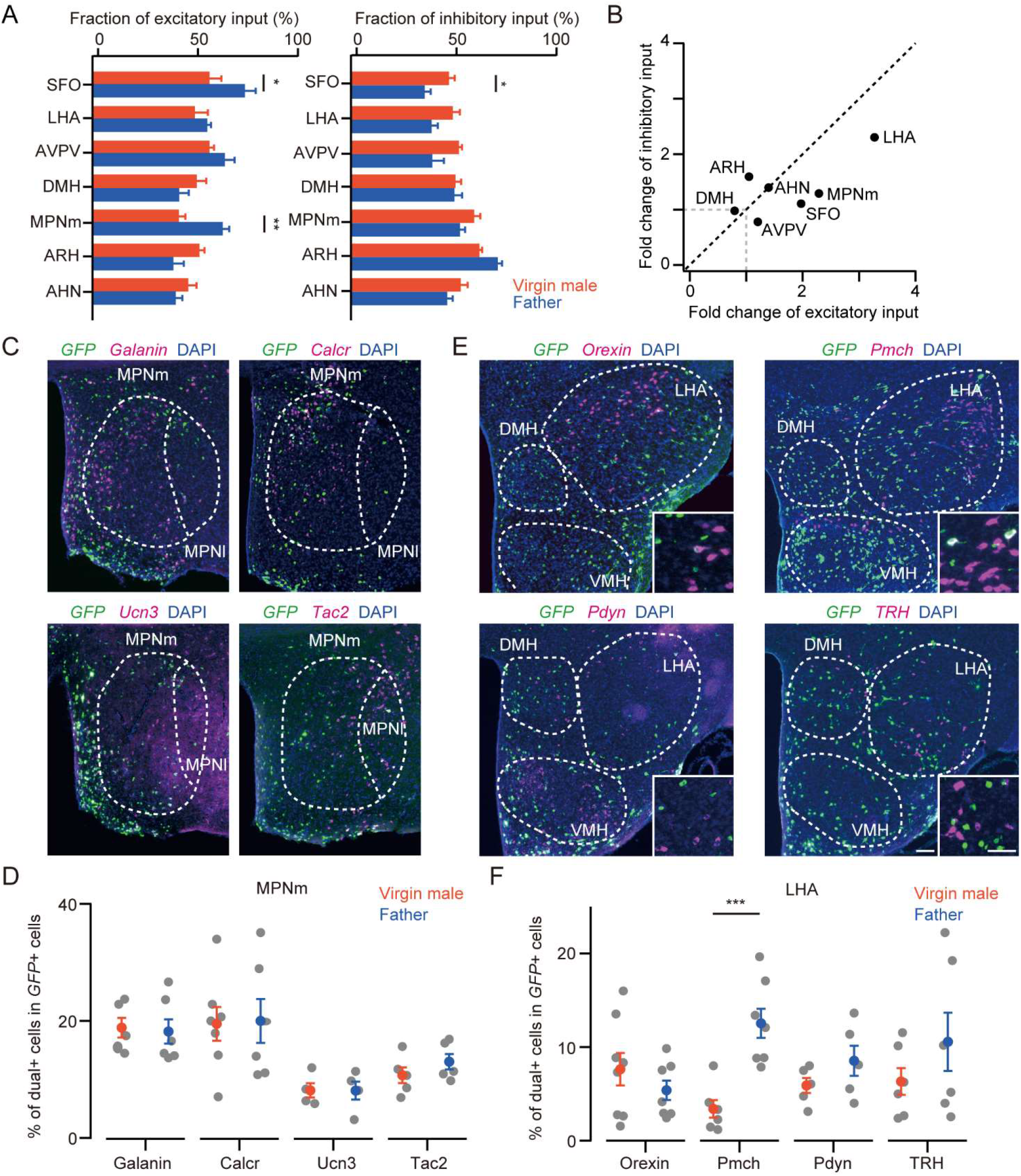
Cell type characterization for presynaptic neurons of OT neurons. (A) Fraction of excitatory (left) or inhibitory (right) inputs from each nucleus, defined by co-labeling of GFP and *vGluT2* or *vGAT* mRNA signals, respectively. n = 6 and 6 for virgin males and fathers, respectively. Fraction of excitatory input, *p = 0.0424, **p = 0.0011; fraction of inhibitory input, *p = 0.0403; two-tailed Welch’s *t*-test. (B) Fold change of excitatory (x-axis) and inhibitory (y-axis) inputs from virgin males to fathers, estimated by the product of convergence index (Figure 4E) and excitatory/inhibitory ratio (A). (C, E) Representative coronal sections of GFP expression by rabies virus (green) and four major cell types in the MPNm (C) and LHA (E) of fathers (magenta). Blue, DAPI. Scale bar, 50 μm. (D) The composition of cell types in the presynaptic inputs from MPNm was similar between virgin males and fathers. The sample sizes in virgin males were *Galanin*, 5, *Calcr*, 7, *Ucn3*, 4, and *Tac2*, 5; in fathers, *Galanin*, 6, *Calcr*, 6, *Ucn3*, 4, and *Tac2*, 5. (F) The ratio of input from LHA *Pmch+* cells was higher in fathers than in virgin males (***p < 0.001, two-tailed Welch’s *t*-test with Bonferroni correction). The sample sizes in virgin males were *Orexin*, 8, *Pmch*, 6, *Pdny*, 5, and *TRH*, 6; in fathers, *Orexin*, 7, *Pmch*, 7, *Pdny*, 5, and *TRH*, 6. Error bars, SEM.

### Validation of enhanced connectivity with electrophysiology

To examine whether the increase in excitatory inputs to OT neurons accompanies enhanced excitatory synaptic transmission, we performed channelrhodopsin 2 (ChR2)-assisted circuit mapping (CRACM)(Petreanu et al., 2007) in acute slice preparations in which OT neurons were specifically visualized by *AAV-OTp-mCherry* (91% ± 0.6% of mCherry+ neurons expressed *OT* mRNA, n = 3 mice). The input resistance measured at the soma, membrane time constant, and spontaneous firing rate were not significantly different between virgin males and fathers (Figure S5A–S5C). To analyze the excitatory transmission, we targeted ChR2 in the LHA, MPNm, and dorsomedial hypothalamus (DMH) by injecting *AAV-FLEx-ChR2(H134R)* into *vGluT2-Cre* mice (Figures 6A, S5D, and S5E). A brief blue light pulse (1 ms) evoked an excitatory response with a short latency (1–8 ms) in OT neurons (Figure 6B and S5F), which was abolished by the glutamate receptor antagonist (Figure S5G), suggesting monosynaptic glutamatergic transmission. Thus, the existence of monosynaptic connections to OT neurons revealed by rabies tracing data (Figure 4E and Figure 5B) was confirmed by CRACM.

**Figure 6.**
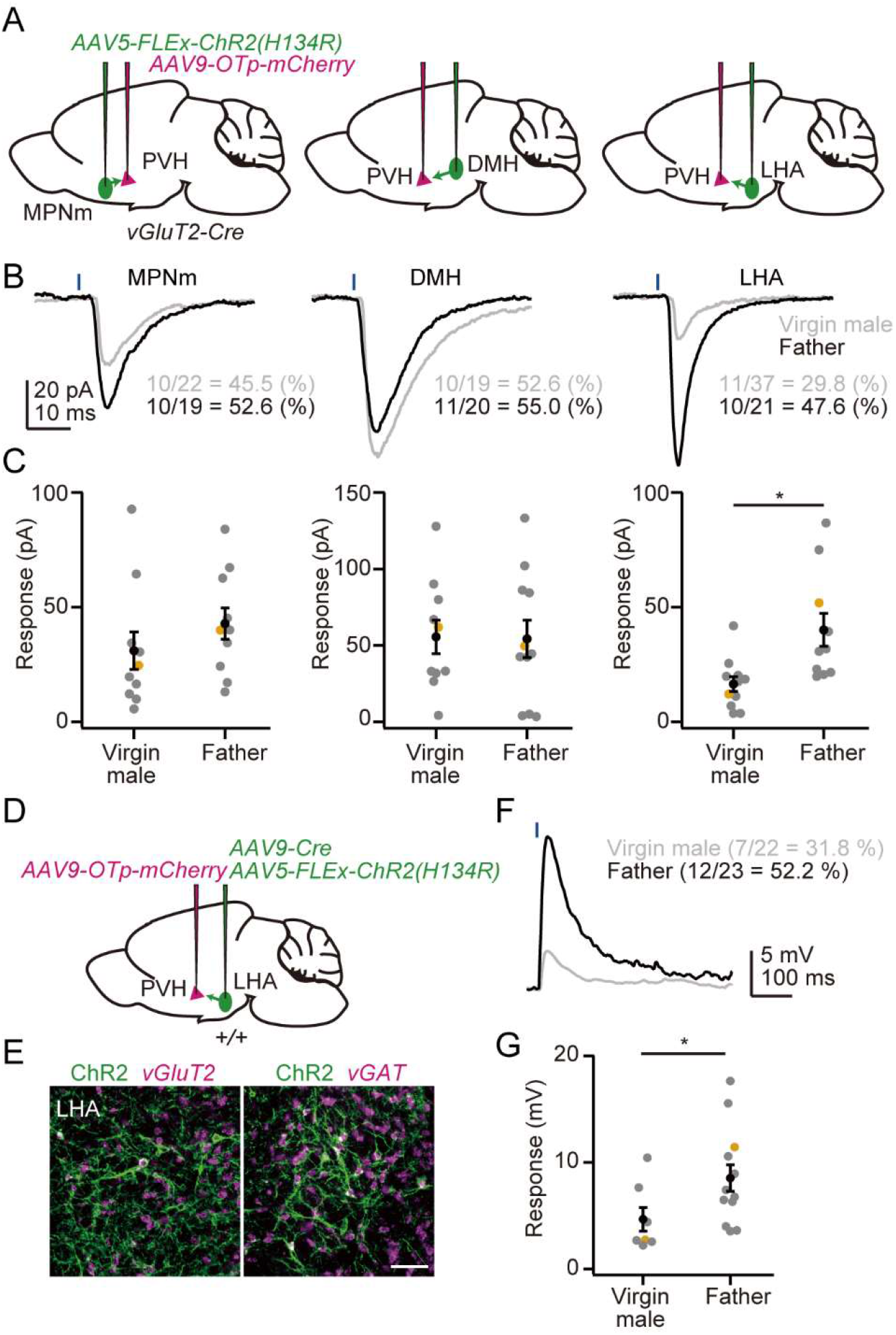
Enhanced excitatory inputs from vGluT2+ LHA neurons to PVH OT neurons by CRACM. (A) Schematic of the virus injections. (B) Representative responses from PVH OT neurons to optogenetic activation of vGluT2+ axons originating from neurons in the MPNm (left), DMH (middle), and LHA (right). Each trace is an average of 10 trials. Blue bar, light stimulation (1 ms). Percentages in each panel indicate the connected ratio. Note that no significance was found in each pair (p > 0.2, two-tailed Fisher’s exact test). (C) The excitatory response was significantly higher in the inputs from LHA (*p = 0.0144, two-tailed Welch’s *t*-test). Orange dots correspond to the traces shown in (B). n = 22 cells from four virgin males and 19 cells from five fathers in the MPNm, 19 cells from four virgin males and 20 cells from five fathers in the DMH, and 37 cells from eight virgin males and 21 cells from six fathers in the LHA. (D) Schematic of the virus injections. (E) Representative coronal sections of the left LHA. Green is ChR2-EYFP amplified by anti-GFP staining. Magenta is *vGluT2* (left) or *vGAT* (right) mRNA *in situ* staining, blue, DAPI. Scale bar, 50 μm. ChR2 was expressed in the *vGluT2*+ and *vGAT*+ cells at a similar ratio in both virgin males and fathers: 54.4% ± 2.8% and 56.0% ± 4.0% of ChR2+ cells co-expressed *vGluT2* in virgin males and fathers (p > 0.79, two-tailed Student’s *t*-test), respectively, and 50.8% ± 1.3% and 50.1% ± 1.3% co-expressed *vGAT* in virgin males and fathers (p > 0.76, two-tailed Student’s *t*-test), respectively. n = 4 for each condition. (F) Representative responses of OT neurons to light stimulation (blue bar, 1 ms). Each trace is an average of 10 trials. Percentages in each panel indicate the connected ratio. (G) Depolarization of OT neurons was larger in fathers (*p = 0.0456. two-tailed Welch’s *t*-test). Orange dots correspond to the traces shown in (F). n = 22 cells from four virgin males and 23 cells from five fathers. Error bars, SEM. See Figure S5 for more data.

Voltage-clamp recordings from OT neurons showed that the excitatory post-synaptic currents evoked by optogenetic stimulation of LHA input were significantly larger in fathers (Figure 6C), consistent with the greatest fold change observed in the trans-synaptic tracing (Figure 5B). Response to MPNm stimulation tended to be enhanced in fathers, but did not reach the level of statistical significance, and response to DMH stimulation was unchanged, which is generally consistent with trans-synaptic tracing. Of note, when we pan-neuronally targeted ChR2 to LHA neurons without restricting to excitatory neurons (Figure 6D and 6E), light stimulation evoked net depolarization in the OT neurons, and the amplitude was still larger in fathers (Figure 6F and 6G). Thus, the overall influence of excitatory and inhibitory circuit changes (Figure 5A and 5B) is biased toward the excitatory drive, consistent with the dominance of excitatory fold change (Figure 5B). Collectively, these results demonstrate enhanced excitatory connectivity from the LHA to OT neurons associated with the life-stage transition.

### Suppression of pup-directed attack by LHA to OT input

LHA has been well-studied in the context of sleep, feeding, stress, and reward (Bonnavion et al., 2016; Petrovich, 2018; Stuber and Wise, 2016). How does the excitatory connection from the LHA to OT neurons contribute to parental behaviors? We first analyzed neural activation of the LHA during pup-directed caregiving behaviors in fathers, based on *c-fos* expression (Figure 7A). We found fathers that interacted with pups (Pup+) expressed *c-fos* at the higher ratio in LHA compared with fathers that were not exposed to pups (Pup−) (Figure 7B and 7C). The majority of *c-fos+* neurons were *vGluT2* positive excitatory neurons (Figure 7D and 7E). Further cell type analysis revealed enhanced *c-fos* expression selectively in the *Pmch*-expressing neurons (Figure 7F and 7G).

**Figure 7.**
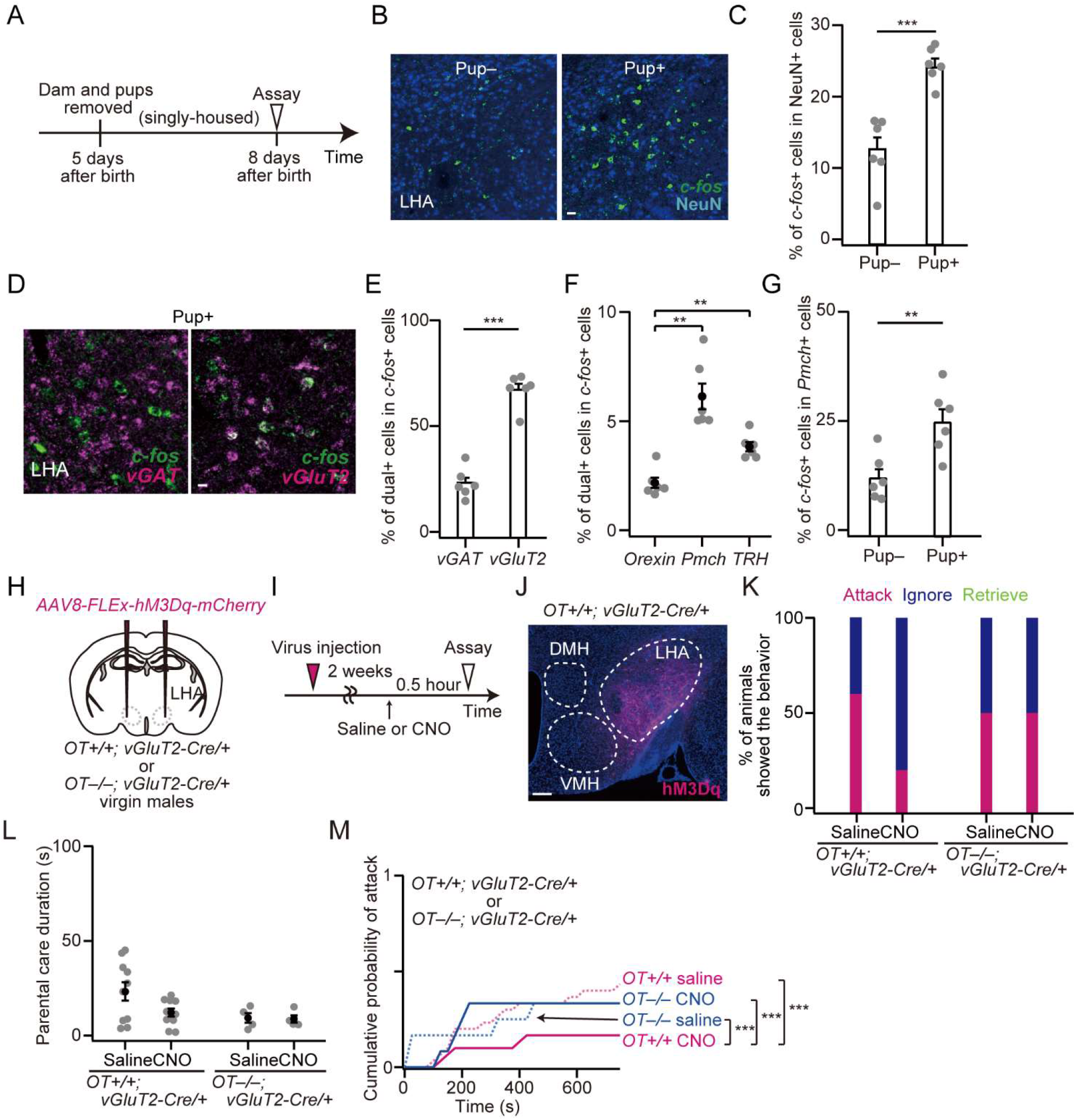
LHA glutamatergic neurons are activated by parental interaction and contribute to the suppression of pup-directed aggression via OT. (A) Schematic of the timeline of the experiment. (B, D) Representative LHA coronal sections from father mice showing *c-fos* mRNA *in situ* staining (green) and anti-NeuN staining (blue) (B) or *c-fos* mRNA (green) and *vGluT2* or *vGAT* mRNA (magenta) *in situ* staining (D). Scale bars, 10 μm. (C, E) Quantification of *c-fos*+ fraction in NeuN+ neurons (C) or a marker positive fraction in *c-fos*+ neurons (E). ***p < 0.001, two-tailed Welch’s *t*-test; n = 6 each. (F) In the three major cell types of excitatory neurons in the LHA, *Pmch+* cells occupy the highest ratio in *c-fos+* activated cells (*Orexin* versus *Pmch*, **p = 0.0091; *Orexin* versus *TRH*, p = 0.0017; one-way ANOVA with repeated measures with post hoc paired *t*-test with Bonferroni correction; n = 6 and 6 for control and parenting, respectively). (G) Enrichment of *c-fos* in *Pmch+* cells was significantly higher in fathers that interacted with pups (**p = 0.0073, two-tailed Welch’s *t*-test; n = 6 and 6 for control and interacted, respectively). (H) Schematic of the virus injections. *AAV-FLEx-hM3Dq-mCherry* was injected into the bilateral LHA of *OT^+/+^; vGluT2-Cre/+* or *OT^−/−^ vGluT2-Cre/+* mice. (I) Schematic of the timeline of the experiment. (J) Representative coronal section around the LHA showing *hM3Dq-mCherry* (magenta) and DAPI (blue). Scale bar, 50 μm. (K) Percentage of animals showing attack, ignore, or retrieve (*OT*^+/+^; *vGluT2-Cre/+*, n = 10 each, and *OT*^−/−^; *vGluT2-Cre/+*, n = 4 each). No pup retrieval was induced by CNO. (L) The duration of parental care was not significantly different (p > 0.05, one-way ANOVA). (M) Cumulative probability of attack to pups. ***p < 0.001, Kolmogorov–Smirnov test with Bonferroni correction. Error bars, SEM. See Figure S6 for more data.

Next, to examine the functional role of excitatory LHA to OT neuron connections in parental behaviors, we targeted *hM3Dq-mCherry* in the *vGluT2+* LHA neurons of *OT*^+/+^ and *OT^−/−^* virgin males (Figure 7H and 7I). In both genotypes, approximately 70% of mCherry+ cells were restricted in the LHA (Figures 7J and S6A). Although CNO injection did not evoke caregiving behaviors (Figures 7K, 7L, and S6B–S6E), we found that *OT*^+/+^ virgin males that received CNO injection were significantly less aggressive toward pups (Figure 7K and 7M). Notably, this suppression of infanticide required functional *OT* gene, as CNO-injected *OT*^−/−^virgin males did not change behaviors (Figure 7M). Therefore, OT release mediated by excitatory LHA neurons contributes to the parental behaviors of fathers via suppressing infanticide.

## Discussion

In the last several decades, the biological mechanisms by which male animals are engaged in direct nurturing behaviors to infants have evoked more general attention, as the caregiving of human fathers to their children has greatly increased across the globe (Feldman et al., 2019). However, basic neuroscience research on paternal caregiving behaviors remains immature, despite their importance. Here we discuss the new insights of neural systems for paternal caregiving behaviors, with respect to the plasticity of neural connections.

### OT neurons facilitate parental behavior in male mice

Our data establish a crucial role of OT neurons and OT ligands for paternal caregiving behaviors. Conditional KO (Figure 1) and whole-body KO (Figure S1) fathers are impaired in the execution of pup-retrieval. These mutant phenotypes in male mice are much clearer than female mice, in which loss of OT or OTR shows subtle phenotype (Macbeth et al., 2010) that may emerge only in stressful environments (Yoshihara et al., 2018). This suggests multiple redundant systems for maternal caregiving behaviors that can compensate for the loss of OT signals. In contrast, male animals are more vulnerable: their pup-directed caregiving behaviors are completely impaired by just a loss of OT (Figure 1) or a loss of prolactin signaling in the MPNm (Stagkourakis et al., 2020). This echoes the fact that even closely related species can be evolved into the opposite ends of the parental-infanticidal spectrum (Bendesky et al., 2017).

The forced activation of OT neurons can confer a full set of parental behaviors such as grooming, crouching, and retrieving, to otherwise infanticidal virgin male mice (Figure 2). How is this effect achieved? We show that activation of OT neurons modulates activities of multiple different cell types: infanticidal PeFA *Ucn3*+ neurons (Autry et al., 2021) are suppressed while parental MPNm *Calcr*+ neurons are activated, when the virgin male mice encountered pups (Figure 2). Previous studies of maternal behaviors showed that auditory (Carcea et al., 2021; Marlin et al., 2015; Schiavo et al., 2020) and olfactory (Oettl et al., 2016) sensory systems are under the influence of OT. Our data add the limbic neural populations directly related to infanticide and parental behaviors to the growing list of neurons that are modulated by OT (Froemke and Carcea, 2017). Of note, another infanticidal cell type, MPN-projecting AHi neurons, can be suppressed by OT via OTR-expressing local interneurons (Sato et al., 2020), suggesting the way of OT action. Future studies will reveal how OTR in various brain regions (Newmaster et al., 2020) adjust circuit functions suited for parental behaviors.

Our data and those from other studies generally suggest a two-step model for inducing paternal caregiving behaviors: Step I to suppress infanticide (ignoring) and Step II to induce a full set of caregiving behaviors (parenting). Step I, but not Step II, can be achieved by gain-of-function of MPNm *galanin+* or *Calcr*+ neurons (Wu et al., 2014; Yoshihara et al., 2021a), loss-of-function of PeFA *Ucn3+* or amygdalo-hippocampal area (AHi) neurons (Autry et al., 2021; Sato et al., 2020), and chemogenetic activation of OT neurons in *OT*-KO background (Figure 3) and *vGluT2+* LHA neurons (Figure 7). In sharp contrast, the OT ligand is indispensable for Step II (Figure 3), which demonstrates the crucial role of OT for paternal caregiving. These gain-of-function effects of OT seem more intensive compared with the recently characterized prolactin system (Stagkourakis et al., 2020). This function of OT may be conserved in humans, as a positive correlation in the plasma OT level and intensity of father–infant interactions was found in human fathers (Gordon et al., 2010).

### Rearrangement of neural connections from virgin males to fathers

The rapid progress of virus-based circuit mapping tools in mice allows comprehensive mapping of input/output wirings of specific neuronal types at the scale of the entire brain (Callaway and Luo, 2015; Oh et al., 2014; Winnubst et al., 2019). However, as such brain-wide connections are often described in a specific sex and reproductive stage (i.e., sexually naïve young-adult males), dynamics of neuronal connections in life-stage transition remain elusive. The present study expands the scope of rabies-based unbiased screening of experience-triggered circuit-level changes (Beier et al., 2017) and demonstrates its utility for identifying naturally-occuring, life-stage-associated changes of connection diagrams without prior assumptions regarding the functional roles of target neurons.

Besides identifying *vGluT2+* LHA neurons as prominently enhanced connections in fathers, our data also suggest widespread and nuanced adjustments (<1.5-fold change) of connections throughout the input nodes (Figure 4E). Although it is technically challenging to validate the function of each minor change, the accumulation of small increases and decreases of connections across diverse brain regions can substantially modify the behavioral output. Indeed, we noticed that the behavioral phenotype obtained by the activation of *vGluT2+* LHA neurons (Figure 7K), or the recently reported activation of LHA MCH neurons (Kato et al., 2021), is much weaker than the activation of post-synaptic OT neurons, suggesting that other connections may be necessary to evoke sufficient activities of OT neurons.

How do neural connections to OT neurons alter upon the life-stage transition? Are they regulated by NMDA-mediated mechanisms known for cortical synaptic plasticity (Nicoll, 2017) and/or sex steroid hormones known for parental functions (Kohl and Dulac, 2018)? Taken together with a remarkable recent study reporting estrus-cycle-related periodic remodeling of a specific hypothalamic connection in virgin female mice (Inoue et al., 2019), our study demonstrated an unexpected degree of structural plasticity in the adult hypothalamus, the brain region classically assumed to be hard-wired. The approach described here can be broadly applied to illuminate neural circuit plasticity in various forms of life-stage transitions.

## STAR☆METHODS

KEY RESOURCES TABLE.

KEY RESOURCES TABLE
| REAGENT or RESOURCE | SOURCE | IDENTIFIER |
| --- | --- | --- |
| **Antibodies** |  |  |
| horseradish peroxidase (HRP) conjugated anti-Dig antibody | Roche Applied Science | cat#11207733910 |
| rat anti-RFP | Chromotek | cat#5f8 |
| mouse anti-Myc | Santa Cruz | cat#sc-40 |
| chicken anti-GFP antibody | Aves Labs | cat#GFP-1010 |
| donkey anti-rat IgG, Cy3 conjugated | Jackson Immuno Research | cat#712-165-153 |
| goat anti-chicken IgY, Alexa Fluor 488 | Jackson Immuno Research | cat#703-545-155 |
| donkey anti-mouse Alexa Fluor 488 | Invitrogen | cat#A32766 |
| **Bacterial and virus strains** |  |  |
| AAV serotype 5 *CAG-FLEX-TVA-mCherry* | Miyamichi et al., 2013 (Custom made by UNC Vector Core) | N/A |
| AAV serotype 8 *CAG-FLEX-RG* | Miyamichi et al., 2013 (Custom made by UNC Vector Core) | N/A |
| AAV serotype 2 *CAG-FLEX-TVA<sup>66T</sup>-mCherry* | Miyamichi et al., 2013 (Custom made by UNC Vector Core) | N/A |
| AAV serotype 2 *CAG-FLEX-RG* | Miyamichi et al., 2013 (Custom made by UNC Vector Core) | N/A |
| AAV serotype 9 *hSyn-Cre* | Addgene | #105555 |
| AAV serotype 8 *hSyn-FLEX-hM3Dq-mCherry* | Addgene | #44361 |
| AAV serotype 5 *EF1a-FLEX-hChR2(H134R)-EYFP* | Addgene | #20298 |
| AAV serotype 8 *hSyn-FLEX-mCherry* | Addgene | #50459 |
| AAV serotype 5 *OTp-hM3Dq-Myc* | This paper (Custom made by UNC Vector Core by using pAAV *OTp-hM3Dq-myc*) | N/A |
| AAV serotype 9 *OTp-mCherry* | This paper (Custom made by Gunma Univ. Vector Core by using pAAV *OTp-mCherry*) | N/A |
| Rabies *dG-GFP+EnvA* | Reconstructed and amplified in the lab | Osakada and Callaway, 2013 |
| **Experimental models: Cell lines** |  |  |
| B7GG cell line | Gift from Dr. Edward Callaway | Osakada and Callaway, 2013 |
| BHK-EnvA cell line | Gift from Dr. Edward Callaway | Osakada and Callaway, 2013 |
| HEK293-TVA800 cell line | Gift from Dr. Edward Callaway | Osakada and Callaway, 2013 |
| **Experimental models: Organisms/strains** |  |  |
| Mouse: *OT-Cre* (Oxytocin-Ires Cre) | JAX Laboratories | Jax#024234 |
| Mouse: *vGluT2-Cre* (*Slc17a6-ires-Cre*) | JAX Laboratories | Jax#028863 |
| Mouse: *OT* KO | This paper | #CDB0204E |
| Mouse: *OT* cKO (floxed) | This paper | #CDB0116E |
| Oligonucleotides |  |  |
| Primers for RNA probes, see Table S6 | This paper (Custom made by Thermo Fisher Scientific) | N/A |
| Recombinant DNA |  |  |
| Plasmid: *pAAV CAG-FLEX-TVA-mCherry* | Miyamichi et al., 2013 | Addgene Plasmid #48332 |
| Plasmid: *pAAV CAG-FLEX-RG* | Miyamichi et al., 2013 | Addgene Plasmid #48333 |
| Plasmid: *pAAV CAG-FLEX-TVA<sup>66T</sup>-mCherry* | Miyamichi et al., 2013 | Addgene Plasmid #48331 |
| Plasmid: *pAAV OTp-hM3Dq-myc* | This paper | N/A |
| Plasmid: *pAAV OTp-mCherry* | This paper | N/A |
| mouse whole brain cDNA | GenoStaff | cat#MD01 |
| Software and algorithms |  |  |
| Igor Pro | Wavemetrics (https://www.wavemetrics.com/) | N/A |
| ImageJ | NIH (https://imagej.nih.gov/ij/) | N/A |
| Other |  |  |
| Linear slicer | Dosaka EM | Pro 7N |
| Single channel heater controller | Warner instruments | TC-324B |

## LEAD CONTACT AND MATERIALS AVAILABILITY

All data are available in the main paper and the supplementary materials. All materials, including the *OT* KO and cKO mice, plasmids, and custom analysis codes, are available through requests to the Lead Contact, Kazunari Miyamichi at.

## EXPERIMENTAL MODEL AND SUBJECT DETAILS

Experimental protocols utilizing rabies virus followed Biosafety Level 2 (P2/P2A) procedures approved by the biosafety committee of the RIKEN Center for Biosystems Dynamics Research (BDR). All animal procedures followed animal care guidelines approved by the Institutional Animal Care and Use Committee of the RIKEN Kobe branch. Wild-type C57BL/6J mice were purchased from Japan SLC. *OT-Cre* (*Oxytocin-ires-Cre*, Jax #024234) and *vGluT2-Cre* (also known as *Slc17a6-ires-Cre*, Jax #028863) mice were purchased from the Jackson Laboratory. Animals were housed under a regular 12-h dark/light cycle with *ad libitum* access to food and water.

## METHOD DETAILS

### Generation of *OT* KO/cKO mice

*OT* KO (Accession. No. CDB0204E) and cKO (Accession. No. CDB0116E) lines (listed in http://www2.clst.riken.jp/arg/mutant%20mice%20list.html) were generated by CRISPR/Cas9-mediated knockin in zygotes, as previously described (Abe et al., 2020). The donor vector consisted of floxed exon 1 of *OT*, encompassing the initiation methionine codon and the signal peptide sequence, which was generated for CRISPR/Cas9-based precise integration into the target chromosome (PITCh) system (Figure S1A)(Sakuma et al., 2016). The guide RNA (gRNA) sites were designed by using CRISPRdirect (Figure S1B)(Naito et al., 2015) to target upstream and downstream of exon 1. Non-homologous end-joining and microhomology-mediated end-joining resulted in the targeted deletion of exon 1 (KO allele) or knockin of floxed exon 1 (cKO allele). For microinjection, the mixture of three crRNAs (CRISPR RNAs)(50 ng/μl), tracrRNA (trans-activating crRNA) (100 ng/μl), donor vector (10 ng/μl), and Cas9 protein (100 ng/μl), were injected into the pronucleus of C57BL/6 one-cell stage zygote. From 223 zygotes, 45 F_0_ founder mice were obtained, 10 of which were double-loxP-positive as identified by PCR and sequencing. To generate the *OT* KO (Accession. No. CDB0204E) line, the genomic deletions between gRNA1 and gRNA2 sites were also confirmed by PCR and sequencing. 18 F_0_ founder mice had at least one *OT* deletion allele. PCR was performed using the following primers: OT-F 5’-cctccaacccctcccaagtc, OT-R1 5’-atccgaatccggactgtggc, and OT-R2 5’-ggtccttagcatcgccagaacg, as shown in Figure S1A. For the sequence analysis, the PCR products from using the primer OT-F and OT-R1 were subcloned into the pCR Blunt II TOPO vector (Zero Blunt TOPO PCR Cloning Kit, ThermoFisher) and sequenced using M13-Foward and M13-Reverse primers. crRNA 1(5’-GGG CCU GCC UCU AAA CAG CGg uuu uag agc uau gcu guu uug), cRNA 2 (5’-GUU ACC UUC ACG GCG UCA AGg uuu uag agc uau gcu guu uug), PITCh 3 crRNA (5’-GCA UCG UAC GCG UAC GUG UUg uuu uag agc uau gcu guu uug), and tracrRNA (5′-AAA CAG CAU AGC AAG UUA AAA UAA GGC UAG UCC GUU AUC AAC UUG AAA AAG UGG CAC CGA GUC GGU GCU) were purchased from FASMAC (Atsugi, Japan). The germline transmissions of *OT* cKO and KO alleles were confirmed by genotyping of F_1_ mice (Figure S1C). Of note, histochemical analysis by *in situ* hybridization (ISH) (Figure 1C and S1D) also confirmed the successful deletion or insertion of gene cassettes, respectively.

### Viral preparations

We obtained the following AAV vectors from Addgene (titer is shown as genome particles [gp] per ml): AAV serotype 9 *hSyn-Cre* (#105555, 2.3 × 10^13^ gp/ml), AAV serotype 8 *hSyn-FLEx-hM3Dq-mCherry* (#44361, 3.2 × 10^13^ gp/ml), AAV serotype 8 *hSyn-FLEx-mCherry* (#50459, 2.3 × 10^13^ gp/ml), and AAV serotype 5 *EF1a-FLEx-hChR2(H134R)-EYFP* (#20298, 7.7 × 10^12^ gp/ml). We subcloned a 2.6-kb mouse *OT* promoter (*OTp*) orthologous to the rat *OTp* (Knobloch et al., 2012) following PCR by using the C57BL/6J mouse genome as a template and PCR primers 5’-agatgagctggtgagcatgtgaagacatgc and 5’-ggcgatggtgctcagtctgagatccgctgt. The AAV vector that expresses mCherry (serotype 9; 2.9 × 10^13^ gp/ml) under the control of the mouse *OTp* was generated by the Gunma University viral vector core. To construct *pAAV-OTp-hM3Dq-Myc*, we subcloned *hM3Dq* from *pAAV-hSyn-FLEx-hM3Dq-mCherry* as a template obtained from Addgene (#44361). 3x*Myc* obtained from *pBT225_(pCA-tdT3Myc)* (#36873, Addgene) was fused to *hM3Dq*. The AAV serotype 5 *OTp-hM3Dq-Myc* (2.9 × 10^12^ gp/ml) was then generated in the UNC vector core.

The following AAV vectors were generated *de novo* by the UNC vector core using the plasmids, as previously described (Miyamichi et al., 2013): AAV serotype 2 *CAG-FLEx-TVA*^*66T*^-*mCherry* (1.0 × 10^12^ gp/ml), AAV serotype 2 *CAG-FLEx-RG* (2.4 × 10^12^ gp/ml), AAV serotype 5 *CAG-FLEx-TVA-mCherry* (2.4 × 10^13^ gp/ml), and AAV serotype 8 *CAG-FLEx-RG* (1.0 × 10^12^ gp/ml).

### Stereotactic injection

To target AAV into a specific brain region, stereotactic coordinates were defined for each brain region based on the Allen Mouse Brain Atlas (Lein et al., 2007). Mice were anesthetized with 65 mg/kg ketamine (Daiichi Sankyo) and 13 mg/kg xylazine (X1251, Sigma-Aldrich) via intraperitoneal injection and head-fixed to stereotactic equipment (Narishige). The following coordinates were used (in mm from the bregma for anteroposterior [AP], mediolateral [ML], and dorsoventral [DV]): PVH, AP −0.8, ML 0.2, DV 4.5; LHA, AP −1.7, ML 1.0, DV 5.2; MPNm, AP −0.2, ML 0.2, DV 5.2; DMH, AP −1.5, ML 0.2, DV 5.0. The injected volume of AAV was 200 nl unless otherwise mentioned at a speed of 50 nl/min. After viral injection, the animal was returned to the home cage.

### Parental behavior assay

#### Assay for virgin males

For the behavioral assay, 10–12-week-old virgin males were used. Before the assay, male mice were housed individually for 5–7 days. Each cage contained shredded paper on wood chips. The mouse builds its nest with the papers, typically at a corner of the cage. Experiments started at 4–6 h before the beginning of the dark phase and were performed under dim fluorescent light. Each male was tested only once. Three 5–7-day-old C57BL/6J pups that had not been exposed to any adult males prior to the experiment were placed in different corners where the testing male had not built its nest. The introduction of pups marked the beginning of the assay. Males were allowed to interact with the pups freely for 15 min. If any sign of pup-directed aggression was observed, however, the targeted pup was immediately rescued from the cage.

The behavior of the animals was categorized based on the following criterion: animals that retrieved at least one pup to the nest were categorized as ‘Retrieve’ and animals that attacked at least one pup were categorized as ‘Attack’. All the remaining animals were scored as ‘Ignore’. Note that, in our data set, animals that showed retrieval of pup(s) did not attack pup(s), and *vice versa*. Therefore, the summation of the ratio of these three categories was always 100%. The following behaviors were further scored: latency to investigate (time after the introduction of pups to the first investigation: typically sniffing a pup), pup retrieval (picking up a pup with its mouth and carrying it to the nest), grooming (licking a pup), crouching (nursing-like posture over pup[s]), and pup-directed aggression (biting a pup or licking a crying pup aggressively). The duration of animals undergoing either grooming, crouching, or retrieving was scored as parental care duration.

For the behavioral assay with chemogenetic activation, AAV that expresses *hM3Dq* was injected 3 weeks beforehand. At 30 min before the assay, CNO (4930, Tocris) dissolved in saline was intraperitoneally injected to achieve a dose of 1 mg/kg. A saline-only injection was used as a control. Each assay was conducted by two experimenters: experimenter 1 prepared two identical tubes containing either saline or saline with CNO, and experimenter 2, who was blinded to the contents of the tubes, conducted the behavioral assay, immunostaining, and data analysis. See Movie S1 for an *OT-Cre* virgin male expressing hM3Dq performing retrieval. In Figures 1–3, 7, S2, and S6, a linear fit was calculated by using data points of all behavioral categories (attack, ignore, and retrieve).

#### Assay for fathers

The behavioral assay with fathers was conducted by using the same procedure as that for virgin males, with several modifications. An individually housed virgin male (8 weeks old) was paired with a female. The next day, the vaginal plug was checked and only the male-female pairs that successfully formed a plug were used for further experiments. Males cohabited with the mated female until 5 days after the birth of pups. The behavioral assay for the males (fathers) was conducted 5 days after the birth of pups. Females (mothers) and pups were removed from the home cage 6–8 h prior to the assay, leaving only the fathers. Unfamiliar pups (pups unrelated to the resident father), prepared as the assay for the virgin males, were used for the assay.

#### Assay for c-fos expression

The behavioral assay for visualizing *c-fos* expression by ISH was conducted with virgin males (Figure 2) and fathers (Figure 7). The assay was performed in the dark. For virgin males, each male was singly housed for 7 days prior to the testing. Each cage contained a metallic tea strainer. On the test day, three pups were packed into the tea strainer, so that males could sense the pups but were physically inaccessible to attack them (note that we prepared this protection because the experimenter could not rescue pups immediately due to the dark environment). Males were allowed to interact with pups for 20 min. After 20 min, males were sacrificed. In fathers, they were allowed to cohabit with a mated female and spend 5 days with the dam and pups. Five days after the birth of the pups, both the dam and pups were removed from the home cage and the father was left alone for 3 days prior to the testing. On the test day, three unfamiliar pups were introduced into the cage of the “Pup+” group, but not into the cage of the “Pup−” group. Each father was allowed to interact freely with the pups for 20 min. After 20 min, fathers were sacrificed as described in the Histochemistry section.

### Electrophysiology

Virgin males and fathers were prepared as described in the behavioral assay section. For whole-cell patch-clamp recordings, mice were deeply anesthetized with isoflurane. Acute coronal slices (250-μm thick) were prepared using a linear slicer (Pro7N, Dosaka EM) in an ice-cold slicing solution containing (in mM): 230 sucrose, 3 KCl, 1.2 KH_2_PO_4_, 10 MgSO_4_, 10 D-glucose, 26 NaHCO_3_, 2 CaCl_2_, 5 Na ascorbate, and 3 Na pyruvate, bubbled with 95% O_2_/5% CO_2_. Slices were incubated and maintained at room temperature for at least 30 min before recording in the solution containing (mM): 92 NaCl, 20 HEPES, 2.5 KCl, 30 NaHCO_3_, 1.2 NaH_2_PO_4_, 2 CaCl_2_, 2 MgSO_4_, 25 D-glucose, 5 Na ascorbate, and 3 Na pyruvate, bubbled with 95% O_2_/5% CO_2_ (pH ~ 7.3, osmolarity adjusted to 295–305 mOsm). Individual slices were transferred into a submerged experimental chamber and perfused with oxygenated ACSF containing (in mM): 126 NaCl, 26 NaHCO_3_, 2.5 KCl, 1.25 NaH_2_PO_4_, 1 MgSO_4_, 12.5 D-glucose, and 2 CaCl_2_, bubbled with 95% O_2_/5% CO_2_ (pH ~ 7.3, osmolarity adjusted to 295–305 mOsm). The temperature of the bath was monitored and adjusted to 32–340 C (TC-324B, Warner Instruments). Whole-cell patch-clamp recordings were performed using an IR-DIC microscope (BX51WI, Olympus) with a water immersion 40 × objective lens (N.A. 0.8). Patch pipettes having a resistance of 4–6 MΩ were fabricated from thin-wall glass capillaries (1.5 mm o.d./1.12 mm i.d., TW150F-3, World Precision Instruments). The internal patch pipette solution used for voltage-clamp recordings contained the following (in mM): 117 cesium methanesulfonate, 20 HEPES, 0.4 EGTA, 2.8 NaCl, 0.3 Na_3_GTP, 4 MgATP, 10 QX314, 0.1 spermine, and 13 biocytin hydrazide (pH ~ 7.3, osmolarity adjusted to 285–295 mOsm). For the current-clamp experiments, the patch pipettes were filled with a solution containing (in mM): 126 K gluconate, 10 HEPES, 0.3 EGTA, 4 KCl, 10 phosphocreatine disodium salt hydrate, 0.3 Na3GTP, 4 MgATP, and 13 biocytin hydrazide (pH ~ 7.3, osmolarity adjusted to 285–295 mOsm). Electrophysiological recordings were made using a Multiclamp 700B amplifier (Molecular Devices), low-pass filtered at 1 kHz, and digitized at 10 kHz. In voltage-clamp mode, cells were held at −70 mV, and in current-clamp mode, cells were held at around −70 mV by injecting a hyperpolarizing current.

#### ChR2-mediated circuit mapping

In Figure 6A–6C, *AAV8-EF1a-FLEx-ChR2(H134R)-EYFP* (100 nl) was injected into the bilateral LHA, MPNm, or DMH of *vGluT2-Cre* mice. In Figure 6D–G, a 1:1 mixture of *AAV8-EF1a-FLEx-hChR2(H134R)-EYFP* and *AAV9-hSyn-Cre* (100 nl) was injected into the bilateral LHA of wild-type mice. OT neurons were labeled by injecting *AAV9-OTp-mCherry* (200 nl) into the bilateral PVH. Recordings were performed 4 weeks after viral delivery. Optogenetic stimulation (1 ms) was achieved through a 470-nm LED light (M470L3, Thorlabs). The optical intensity of the LED light was adjusted to 3.6 mW measured at the back aperture of the objective lens (S120VC sensor, Thorlabs). OT neurons were targeted by the fluorescence of mCherry, illuminated by a 590-nm LED light (M590L3, Thorlabs).

Optogenetic responses were recorded in voltage-clamp mode (Figure 6A–C) and current-clamp mode (Figure 6F and 6G). From each cell, we obtained 10–15 traces. The recorded OT neuron was regarded as “connected” to the stimulated neurons if the cell showed a time-locked response in 80% or higher trials within 10ms after a light pulse. The response amplitude was calculated by subtracting the baseline (average of 50 ms before light pulse) from the average of 5 ms around the peak. The response onset time was detected by an algorithm described previously (Kudoh and Taguchi, 2002). In Figure S5A–S5C, all physiological properties were measured shortly after attaining the whole-cell configuration. In Figure 6B and 6F, each trace was an average of 10 trials.

### Trans-synaptic retrograde tracing

The preparation of rabies virus was conducted by using viruses and cell lines as previously described(Osakada and Callaway, 2013). RV*d*G-GFP was *de novo* prepared by using the B7GG cell line (a gift from Ed Callaway) and plasmids (*pCAG-B19N*, *pCAG-B19P*, *pCAG-B19L*, *pCAG-B19G*, and *pSADdG-GFP-F2*; gifts from Ed Callaway), and then pseudotyped by using BHK-EnvA cells (a gift from Ed Callaway). The titer of RV*d*G-GFP+EnvA used in this study was estimated to be 4 × 10^9^ infectious particles per ml based on serial dilutions of the virus stock followed by infection of the HEK293-TVA800 cell line (a gift from Ed Callaway).

For trans-synaptic tracing using rabies virus, 200 nl of a 1:1 mixture of AAV5 *CAG-FLEx-TVA-mCherry* and AAV8 *CAG-FLEx-RG* or a mixture of AAV2 *CAG-FLEx-TVA^66T^-mCherry* and AAV2 *CAG-FLEx-RG* (Miyamichi et al., 2013) was injected into the unilateral PVH of *OT-Cre* mice. Two weeks later, 200 nl of RV*dG-GFP*+EnvA was injected into the same brain region to initiate trans*-*synaptic tracing. One week after the injection of rabies virus, mice were sacrificed and perfused with PBS followed by 4% PFA in PBS. 20-μm coronal sections throughout the hypothalamus and anterior part of the thalamus were collected, and every fourth section was subjected to cell counting. In Figure S4, although the experimental conditions of “virgin males” and “5 days after birth” were the same as those for “virgin males” and “Fathers” in Figure 4, respectively, different sets of mice were prepared, providing a different cohort of data. The numbers of GFP+ and mCherry+ cells on the injected hemisphere were counted and assigned to brain areas based on the classification of Allen Mouse Brain Atlas (Lein et al., 2007). Cells were counted manually using the ImageJ Cell Counter plugin. In Figure S4D–S4F, S4I, and S4J, cells were counted by a different experimenter, who was blinded to the experimental conditions, from the experimenter who analyzed Figure 4. Cells that expressed both GFP and mCherry from AAV *CAG-FLEx-TVA-mCherry* or *CAG-FLEx-TVA^66T^-mCherry* were counted as starter cells (Figure 4B and S4D). In the experiments reported in Figure 4, 97.9% ± 1.5% and 99.4% ± 0.52% of starter cells were located in the PVH of virgin males and fathers, respectively. Because we collected every fourth section, the reported number of starter cells (Figure 4B and S4D) was compensated (×4) from the measured value. In Figure 4E and S4J, the d′ value was calculated as

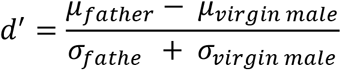

where μ and σ denote the average and standard deviation, respectively. A d′ value larger than 0 represents a single PVH OT neuron in fathers receiving denser input from the nucleus (Figure 4E) or cell type (Figure S4J) compared with virgin males. In Figure S4F, the d′ value was calculated as above, but “father” should be replaced with the three conditions other than “virgin male”.

### Histochemistry

Mice were anesthetized with sodium pentobarbital and perfused with PBS followed by 4% PFA in PBS. The brain was post-fixed with 4% PFA overnight. Twenty-micron coronal brain sections (every fourth section) were made using a cryostat (Leica).

Fluorescent ISH was performed as previously described (Ishii et al., 2017). The primer sets used in the present study are shown in Table S6. Immunohistochemistry was performed as previously reported (Ishii et al., 2017). The following chemicals were used for primary antibodies: anti-GFP (GFP-1010, Aves Labs, 1:500), anti-RFP (5f8, Chromotek, 1:500), and anti-Myc (sc-40, Santa Cruz, 1:1000). Signal-positive cells were detected by anti-chicken Alexa Fluor 488 (703-545-155, Jackson Immuno Research, 1:500), anti-rat Cy3 (712-165-153, Jackson Immuno Research, 1:500), and anti-mouse Alexa Fluor 488 (A32766, Invitrogen, 1:500). Fluoromount (K024, Diagnostic BioSystems) was used as a mounting medium. Brain images were acquired using an Olympus BX53 microscope equipped with a 10× (N.A. 0.4) objective lens. Cells were counted manually using the ImageJ Cell Counter plugin.

### Data analysis

All mean values are reported as mean ± SEM. The statistical details of each experiment, including the statistical tests used, the exact value of n, and what n represents, are shown in each figure legend. The p-values are shown in each figure legend or panel; non-significant values are not noted.

**Table S1. List of abbreviations, related to Figure 4.**

**Table S2. Complete data set of input fraction, related to Figure 4.**

**Table S3. Complete data set of convergence index, related to Figure 4.**

**Table S4. Complete data set of input fraction, related to Figure S4.**

**Table S5. Complete data set of convergence index, related to Figure S4.**

**Table S6. List of primers used in this study.**

**Movie S1. An *OT-Cre* virgin male expressing hM3Dq performing pup retrieval after CNO injection.**

## Author contributions

K.I. and K.M. conceived the experiments. K.I. and M.H. performed the experiments and analyzed the data. K.T. and K.M. generated pseudotyped rabies virus. K.I., M.H., and K.M. performed construction of AAVs, and A.K. and H.H. generated ready-to-use AAV vectors. T.A. and H.K. generated genetically engineered mice. K.I. and K.M. designed the experiments and wrote the paper.

## Acknowledgments

We thank Liqun Luo, Hokto Kazama, and members of the Miyamichi lab for the critical reading of the manuscript, Edward M Callaway for sharing B7GG, BHK-EnvA, and HEK293-TVA800 cell lines, the UNC vector core for the AAV production, Fumitaka Kimura for the advice on electrophysiology, and Masayo Takahashi, Michiko Mandai, and Minoru Takasato for sharing equipment. This work was supported by the RIKEN Special Postdoctoral Researchers Program, a grant from the Kao Foundation for Arts and Sciences, and JSPS KAKENHI (19J00403 and 19K16303) to K.I., JSPS KAKENHI (19K06899) to A.K., the Brain/MINDS program from AMED (JP20dm0207057 and JP21dm0207111) to H.H., and the JST PRESTO program (JPMJPR1789), the Uehara Memorial Foundation Research Grant, a Takeda Science Foundation Research Grant, and JSPS KAKENHI (18H02548 and 20K20589) to K.M.

## Supplementary Figures

**Figure S1.**
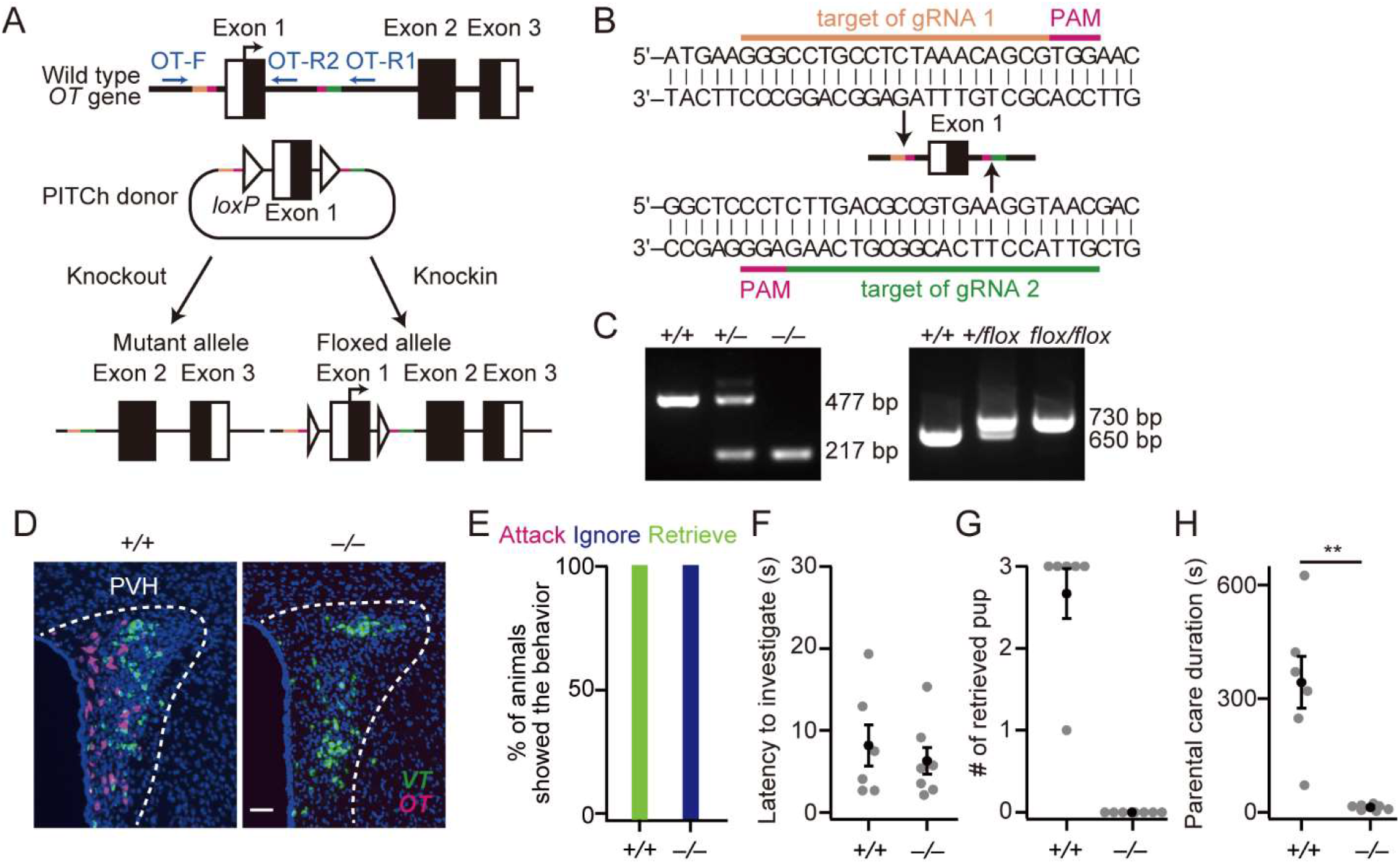
Generation of *OT* KO/cKO mice and behavioral performance of *OT* KO fathers, related to Figure 1. (A) Schematic showing the procedure used to generate *OT* KO and cKO mice using the PITCh system. Top: The *OT* gene consists of three exons (exons 1–3). Open and closed boxes denote untranslated and protein-coding regions, respectively. Blue arrows indicate the site of genotyping primers. Pink line indicates the protospacer adjacent motif (PAM) sequence. Orange and green lines indicate CRISPR target sequences, designed to target upstream and downstream of exon 1 of *OT*, which includes the initiation *ATG* codon and the OT peptide sequence. Middle: Schematic showing the construct of the PITCh donor vector. White triangles denote *loxP*. Bottom: Non-homologous end-joining and microhomology-mediated end-joining result in the targeted deletion of exon 1 (KO) or knockin of loxP sites (cKO). (B) Schematic showing the target sequence of guide RNAs (gRNAs). (C) Representative results of gel electrophoresis examining the KO allele (left) and the floxed allele (right). (D) Representative coronal sections showing *VT* mRNA (green) and *OT* mRNA (magenta) in the PVH of *OT*^+/+^ male mouse (top) and *OT^−/−^* (bottom). The *OT* mRNA signal was undetectable in *OT^−/−^*, whereas the *VT* mRNA signal persisted. Blue, DAPI. Scale bar, 50 μm. (E) Percentage of fathers showing attack, ignore, or retrieve. (F) Latency to the first investigation of pups was not significantly different between *OT^−/−^* and *OT*^+/+^ (p > 0.1, two-tailed Welch’s *t*-test). (G) Number of retrieved pups. (H) Duration of parental care of *OT^−/−^* fathers was significantly shorter than that of *OT*^+/+^ fathers (**p = 0.0071, two-tailed Welch’s *t*-test). n = 6 and 7 for and *OT*^+/+^ and *OT^−/−^*mice, respectively. Error bars, SEM.

**Figure S2.**
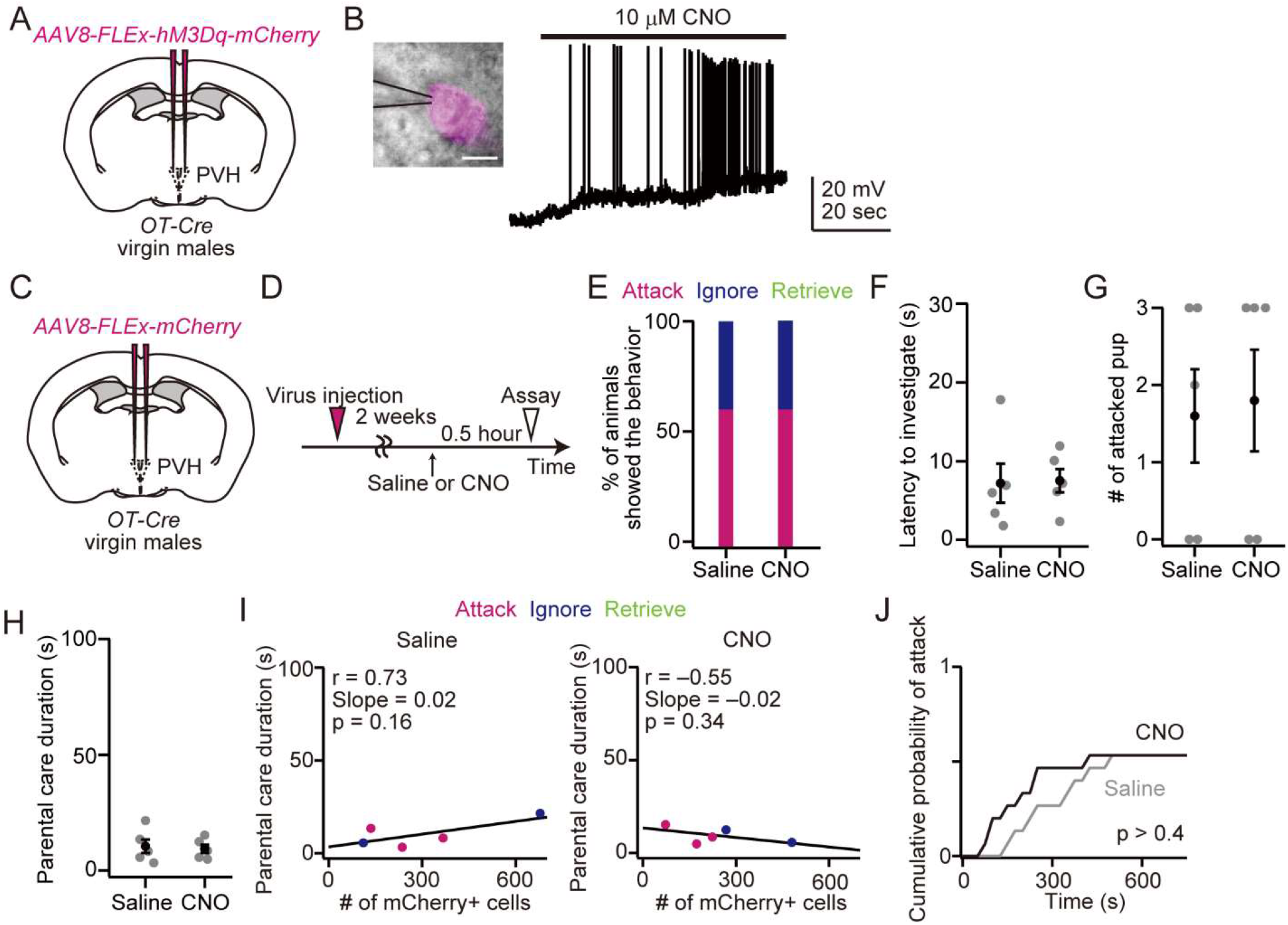
Control experiments for the chemogenetic activation of OT neurons in virgin males, related to Figure 2. (A) Schematic of the virus injection. *AAV-FLEx-hM3Dq-mCherry* was injected into the bilateral PVH of virgin males. (B) Left: whole-cell patch-clamp recording from a representative hM3Dq+ OT neuron in the PVH. Scale bar, 10 μm. Right: Representative response to 10 μm CNO. The result was replicated in three hM3Dq-expressing neurons from three mice. (C) Schematic of the virus injection. AAV-*FLEx-mCherry* was injected into the bilateral PVH of virgin males. (D) Schematic of the timeline of the experiment. (E) Percentage of animals showing attack, ignore, or retrieve. In the virgin males that expressed only mCherry, CNO did not influence the behavior. n = 5 and 5 for saline and CNO, respectively. (F–H) Quantification of behavioral parameters. Latency to the first investigation of pups (F), number of attacked pups (G), and duration of parental care (H). Error bars, SEM. (I) Relationship between parental interaction and the number of mCherry+ neurons in the PVH. Animals are color-coded by their behavioral categories. Black line, a linear fit. (J) Cumulative probability of attack to pups. The p-value is shown in the panel (Kolmogorov– Smirnov test).

**Figure S3.**
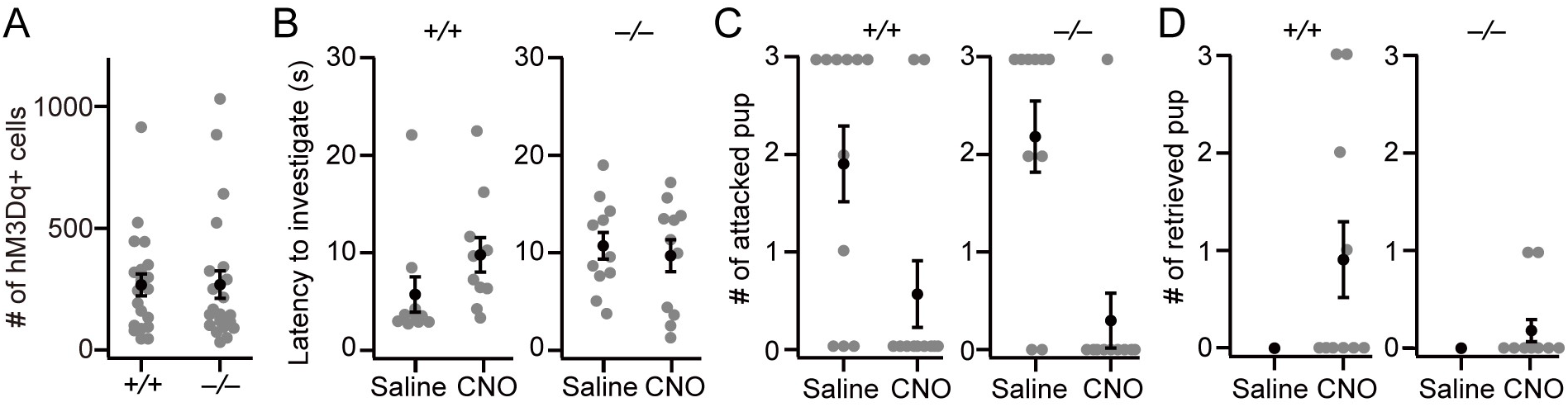
Behavioral performance of *OT*^+/+^ and *OT*^−/−^ virgin males, related to Figure 3. (A) The number of hM3Dq-Myc+ cells was not significantly different between *OT^−/−^* and *OT*^+/+^ (p > 0.5, two-tailed Welch’s *t*-test). (B) Latency to the first investigation of pups. (C) Number of attacked pups. (D) Number of retrieved pups. *OT*^+/+^, n = 10 each, and *OT*^−/−^, n = 11 each, for saline and CNO. Error bars, SEM.

**Figure S4.**
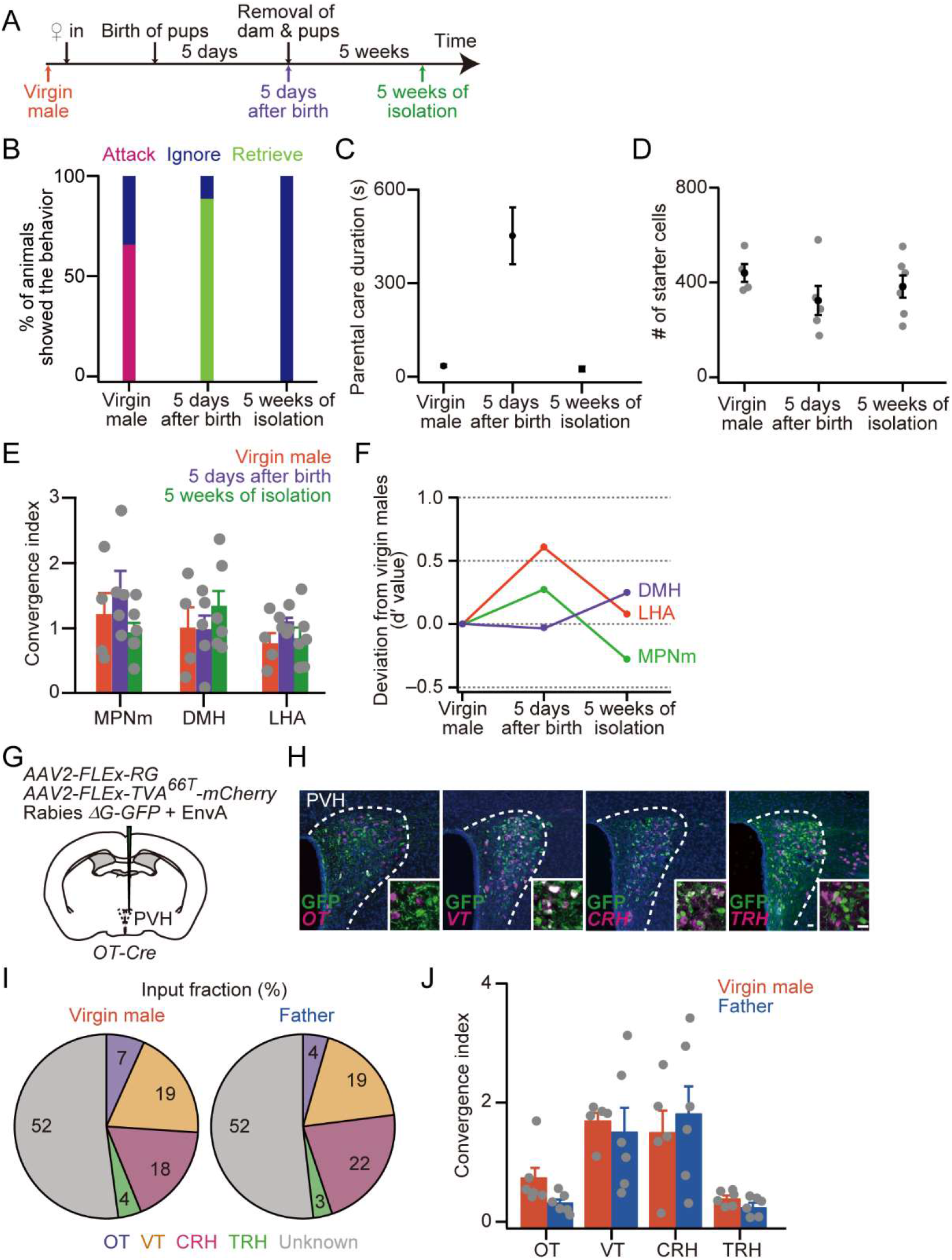
Time course of plasticity in inputs to OT neurons and local neural connections to OT neurons, related to Figure 4. (A) Schematic of the timeline of the experiment. Color-coded arrows indicate the timing of rabies virus injection and behavioral assay. (B) Percentage of males showing attack, ignore, or retrieve. Note that in our strain, fathers with 5 weeks of isolation did not show infanticidal behavior as described previously (vom Saal, 1985). n = 9 each for virgin males, 5 days after birth, and 5 weeks of isolation. (C) Duration of parental care. (D) The number of starter cells was not significantly different (p > 0.4, one-way ANOVA). (E, F) Convergence index (E) and d′ value (F), calculated between each condition and virgin males. Note that the trans-synaptic tracing data presented in this figure are a different cohort of mice and cells counted by a different experimenter, who was blinded to the experimental conditions, from the one who analyzed Figure 4. n = 4, 5, and 6 for virgin males, 5 days after birth, and 5 weeks of isolation, respectively. (G) Schematic showing virus injections. TVA with a point mutation, TVA^66T^, was used to visualize the presynaptic cells within the PVH because of the near-zero Cre-independent transduction of the EnvA-pseudotyped rabies virus. (H) Representative coronal sections of the PVH in virgin males. mRNA of four major cell types in the PVH, *OT*, *VT*, *CRH*, and *TRH*, was visualized by ISH (magenta). Green, GFP stained by anti-GFP, blue, DAPI. Scale bar, 20 μm. (I) Fraction of monosynaptic inputs to OT neurons in virgin males (left) and fathers (right). The input pattern was largely similar. Note that in both virgin males and fathers, the number of inputs from *OT+* neurons may have been underestimated given that a fraction of *OT+* neurons expressed both mCherry (TVA^66T^) and GFP (rabies virus), recognized as starter cells. The number of starter cells in the sections for which we performed *in situ* staining to detect *OT* mRNA was 22.8 ± 4.3 for virgin males and 27.2 ± 4.0 for fathers, occupying 11.9% ± 2.4% and 15.5% ± 2.0% of *OT*+ neurons in those sections in virgin males and fathers, respectively (p > 0.3, two-tailed Welch’s *t*-test). (J) Number of inputs to single OT neurons in virgin males and fathers. Data obtained from six virgin males and six fathers. No significant difference was observed in each pair (p > 0.1, two-tailed Welch’s *t*-test with Bonferroni correction). The d′ values for each cell type were −0.69, −0.15, 0.17, and −0.53 for *OT+*, *VT+*, *CRH+*, and *TRH+*, respectively. See Table S3 and S4 for raw cell counts and sample size.

**Figure S5.**
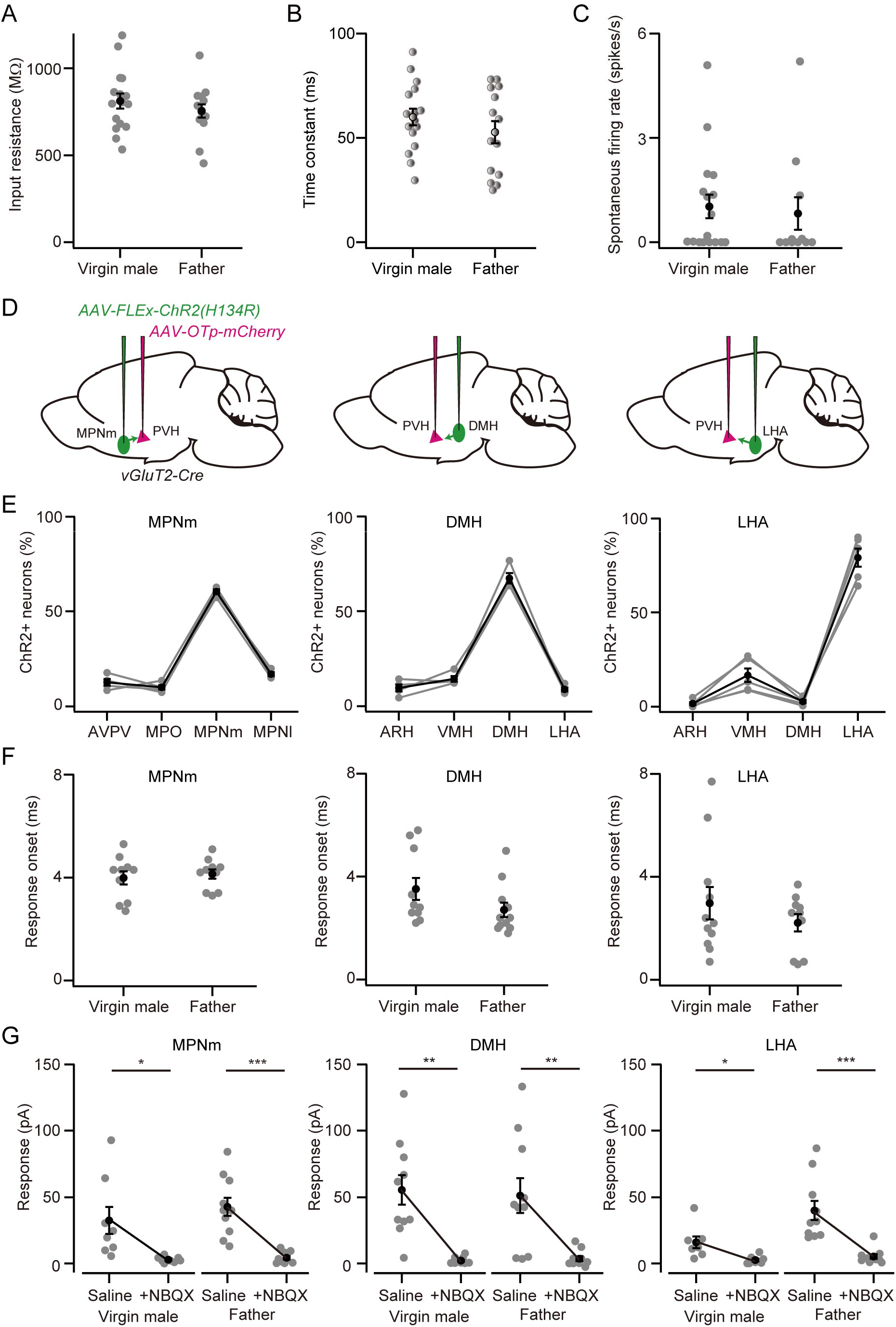
Cellular properties of OT neurons in virgin males and fathers, related to Figure 6. (A, B) Input resistance measured at soma (A) and membrane time constant (B) were not significantly different between virgin males and fathers (n = 16 cells from six virgin males and 14 cells from four fathers; p > 0.3, two-tailed Welch’s *t*-test). (C) The spontaneous firing rate was not significantly different between virgin males and fathers (n = 17 cells from four virgin males and 11 cells from four fathers; p > 0.3, two-tailed Welch’s *t*-test). (D) Schematic of the virus injections. (E) Cells expressing ChR2-EYFP were counted in the targeted nucleus and three neighboring hypothalamic nuclei because *vGluT2-Cre* may express ChR2 other than the targeted area. n = 4, 4, and 5 fathers in the MPNm, DMH, and LHA, respectively. (F) Response onset of data shown in Figure 6C. (G) Optogenetic activation evoked glutamatergic synaptic transmission in both virgin males and fathers, given that application of NBQX significantly decreased the response (MPNm, *p = 0.0245, ***p < 0.001; DMH, **p = 0.0013, **p = 0.0045; LHA, *p = 0.0200, ***p < 0.001; two-tailed paired *t*-test with Bonferroni correction; n = 8 cells from four virgin males and n = 10 cells from five fathers in the MPNm, n = 10 cells from four virgin males and n = 10 cells from five fathers in the DMH, n = 7 cells from six virgin males and n = 10 cells from six fathers in the LHA). Error bars, SEM.

**Figure S6.**
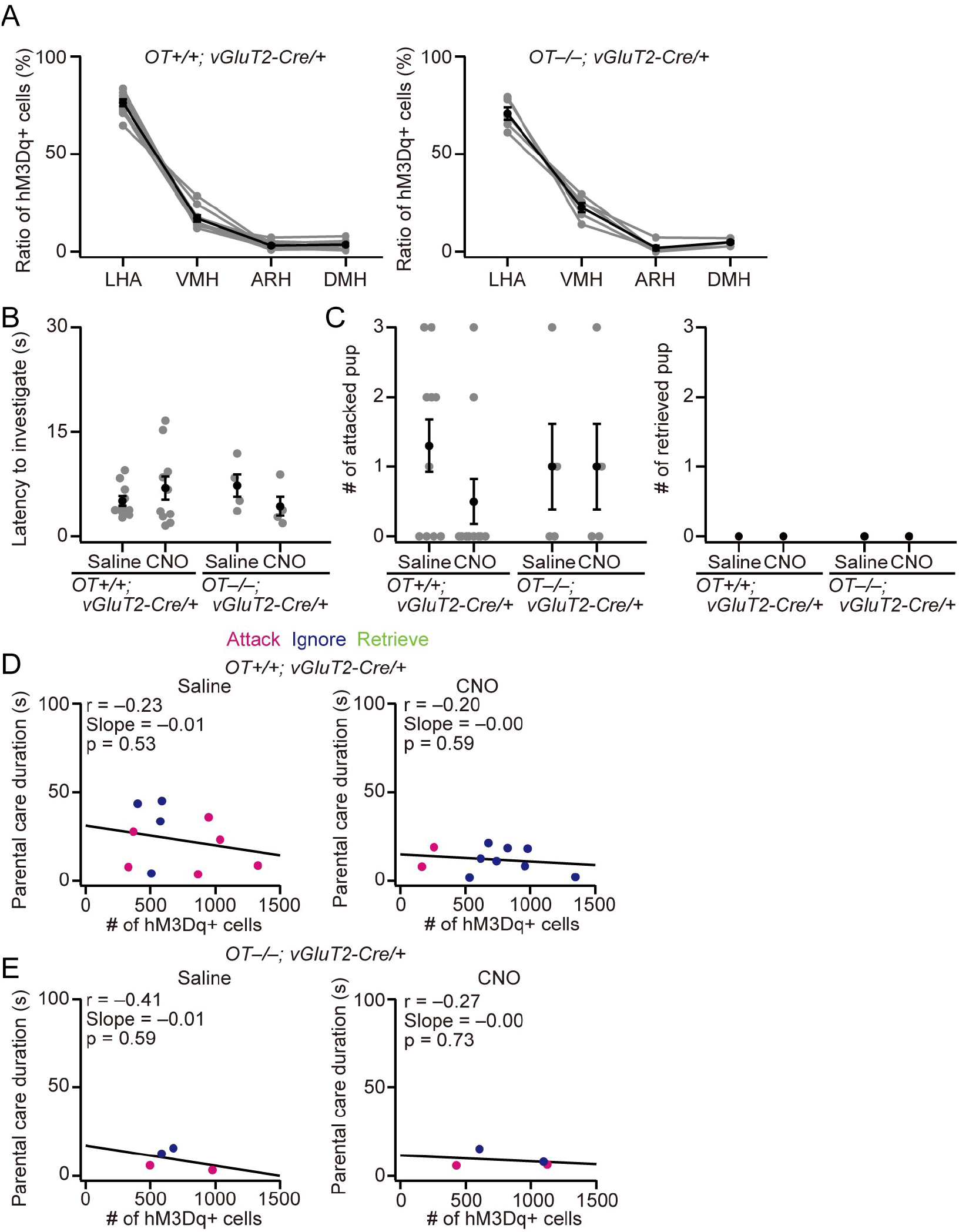
Supplemental information for chemogenetic activation of vGluT2+ neurons in LHA, related to Figure 7. (A) Fraction of neurons expressing hM3Dq in the four neighboring nuclei. n = 10 and 4 for *OT^+/+^; vGluT2-Cre/+* and *OT^−/−^; vGluT2-Cre/+* mice, respectively. (B) Latency to the first investigation of pups was not statistically different. n = 10 each for *OT^+/+^; vGluT2-Cre/+* and n = 4 each for *OT^−/−^; vGluT2-Cre/+*. (C) Numbers of attacked (left) and retrieved (right) pups. (D, E) Relationships between parental interaction and the number of neurons expressing hM3Dq in the LHA of *OT^+/+^; vGluT2-Cre/+* (D) and *OT^−/−^; vGluT2-Cre/+* (E) mice. Animals are color-coded by their behavioral categories. The black line represents a linear fit.

